# Evaluation of a predictive method for the H&E-based molecular profiling of breast cancer with deep learning

**DOI:** 10.1101/2022.01.04.474882

**Authors:** Salim Arslan, Xiusi Li, Julian Schmidt, Julius Hense, Andre Geraldes, Cher Bass, Keelan Brown, Angelica Marcia, Tim Dewhirst, Pahini Pandya, Shikha Singhal, Debapriya Mehrotra, Pandu Raharja-Liu

## Abstract

We present a public validation of PANProfiler (ER, PR, HER2), an in-vitro medical device (IVD) that predicts the qualitative status of estrogen receptor (ER), progesterone receptor (PR) and human epidermal growth factor receptor 2 (HER2) by analysing the hematoxylin and eosin (H&E)-stained tissue scan. In public validation on 648 (ER), 648 (PR) and 560 (HER2) unseen cases with known biomarker status, the device achieves an accuracy of 87% (ER), 83% (PR) and 87% (HER2). The validation offers early evidence of the ability to predict clinically relevant breast biomarkers from an H&E slide in a relevant clinical setting.

## Introduction

Breast cancer is among the most commonly diagnosed cancers, with an estimated 2.3 million new cases worldwide and 685,000 deaths globally in 2020 [1], and an expected increase in the incident cases by more than 46% by 2040 [2]. At the end of 2020, there were 7.8 million women alive who were diagnosed with breast cancer in the past 5 years [3]. Molecular profiling of the estrogen receptor (ER), progesterone receptor (PR) and the human epidermal growth factor receptor 2 (HER2) is performed for all malignant breast cancers to determine the course of cancer development and to inform the choice of targeted therapy for an individual patient. The primary techniques for profiling the molecular biomarkers of breast cancer are immunohistochemistry (IHC) and in-situ hybridization (ISH). These tests pose a great challenge for the current standard practice, as they account for a large proportion of the high turnaround time for breast cancer biopsy diagnosis due to IHC/ISH preparation requiring specialised, quality-assured, laboratory work with dedicated equipment, expensive reagents and additional pathologist time. Sources of delays with the conventional diagnostic workflow and biomarker testing in breast cancer histopathology can be seen in **Figure 1**. These limitations necessitate the development of new tests that can accelerate the cancer diagnosis process, which hold a great potential to reduce pathology workloads, streamline diagnostic workflows, shorten turnaround times and reduce costs.

**Figure 1:**
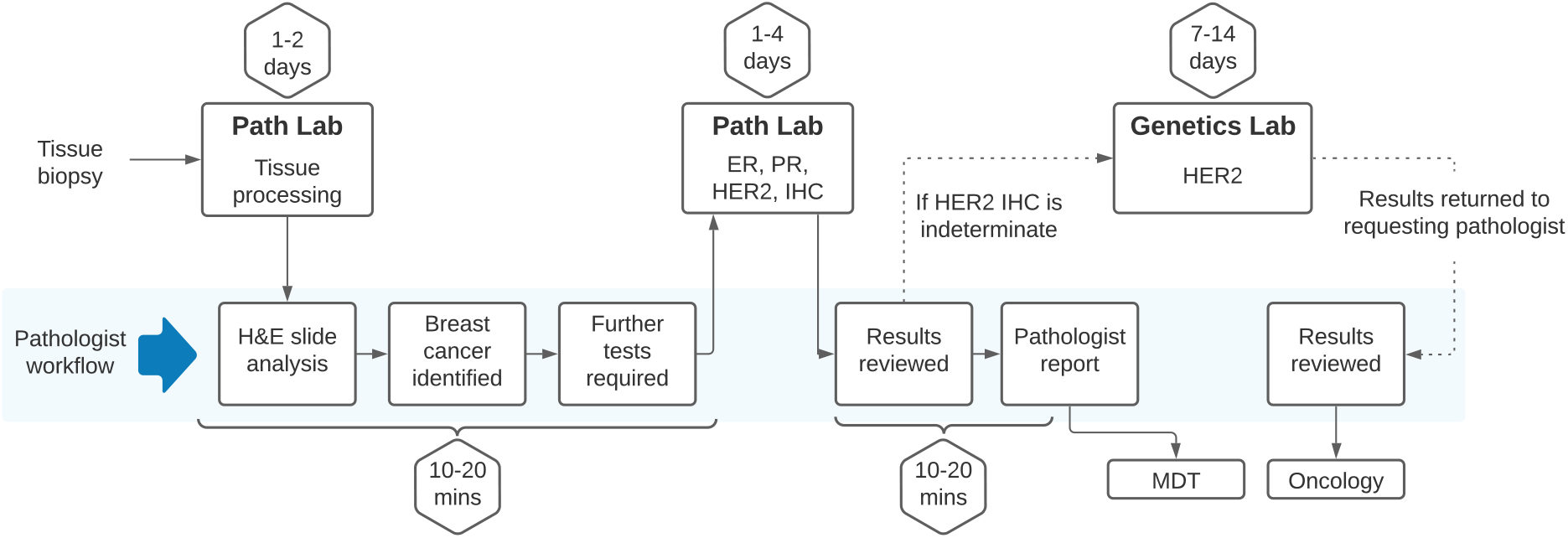
Conventional histopathology workflow for breast cancer diagnosis and biomarker testing. This figure is based on UK-specific primary research i.e. conversations with pathologists and refined based on different settings that our qualified histopathologists have worked in. Much more complex versions of the workflows can be seen in [4]. The time estimates may vary from lab to lab.

Recent advancements in machine learning (ML) and computational pathology have allowed the prediction of certain biomarkers directly from the standard H&E-stained whole slide images, which can potentially reduce the use of standard tests. Accumulating evidence from the literature suggests that H&E images contain information that can be used to infer molecular profiles [5, 6, 7]. Several studies have shown the effectiveness of deep learning (DL) methods, in particular, convolutional neural networks (CNNs), to reveal histopathological patterns in routine H&E images that are correlated with the status of molecular biomarkers in various cancer types, including colorectal [6, 8], lung [5, 9], prostate [10] and skin [11]. Recent pan-cancer studies identified multiple clinically relevant molecular biomarkers across major solid tumours to investigate the links between genetic/molecular alterations and histomorphological features using H&E images [7, 12, 13]. Furthermore, several studies assessed the predictability of ER, PR, HER2 markers in breast cancer from H&E slides [14, 15, 16, 17]. These studies paved a promising pathway to the clinical use-case development of H&E-based biomarker profiling methods for breast cancer.

Building upon the evidence from the scientific literature, we have developed PANprofiler Breast (ER, PR, HER2), a DL-based image analysis software device for the molecular profiling of breast cancer that can analyse visual features from routine histopathological images stained with H&E to predict the status of ER, PR and HER2 receptors. The method combines several steps currently performed in the traditional breast cancer diagnostic pathway into one single pipeline, allowing the prediction of the expression status directly from H&E slides without any intermediate staining and manual quantification. **Figure 2** shows the typical breast cancer diagnostic pathway with traditional molecular testing (red panel) and how the pathway can be improved with the integration of PANProfiler Breast (ER, PR, HER2) (blue panel).

**Figure 2:**
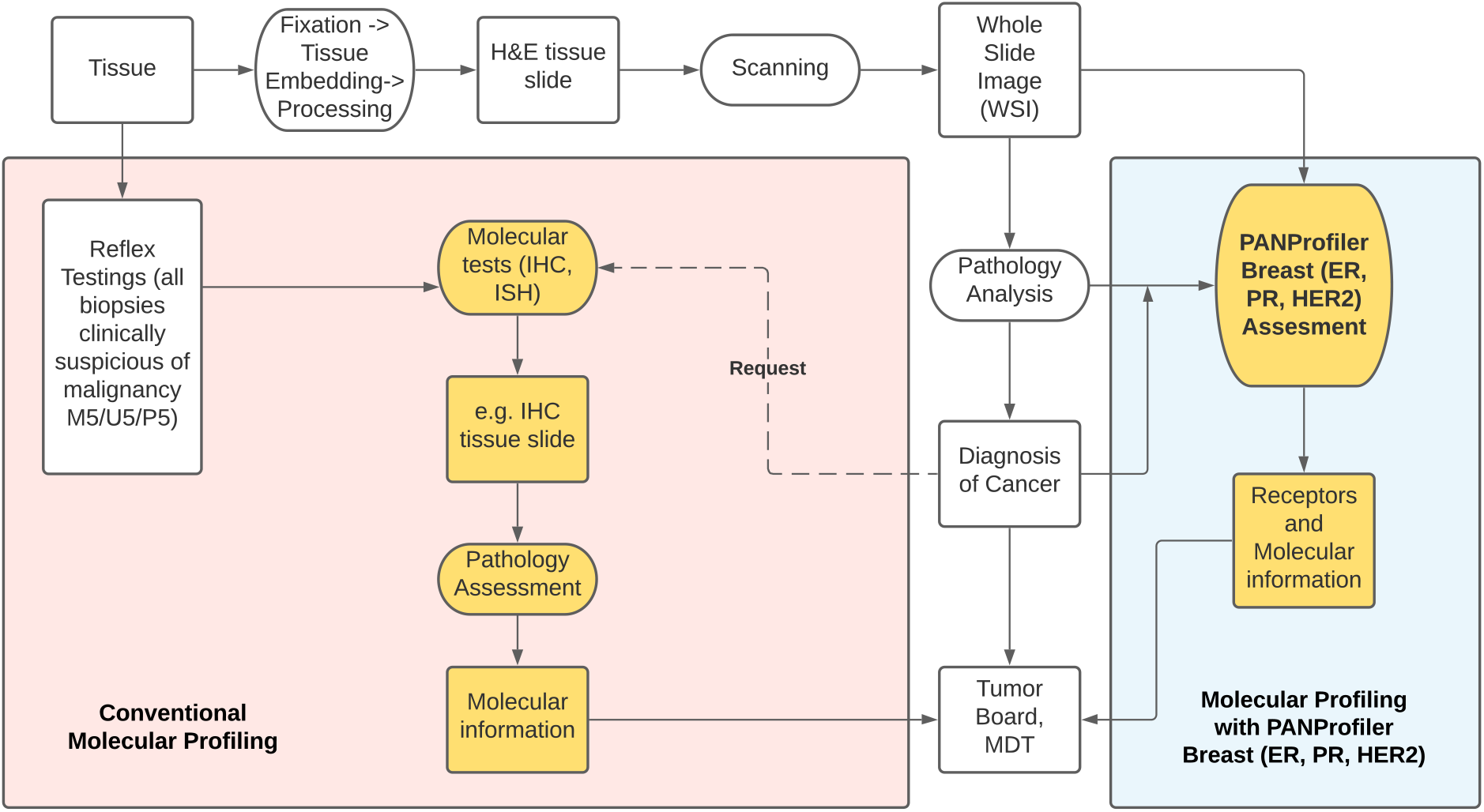
Breast cancer diagnostic pathway with the traditional molecular testing (red panel) and with PANProfiler Breast (ER, PR, HER2) (blue panel).

This paper provides a summary of the approaches and key results acquired from a performance evaluation study carried out to assess the clinical safety and efficacy of PANProfiler Breast (ER, PR, HER2) for its intended purpose. First, a study was carried out to internally validate the effectiveness of the device for each of the biomarkers and to select the predictive models that will constitute the core of the device. A clinical performance evaluation study was then conducted to demonstrate the ability of the device to yield results that are correlated with the well-characterised receptor status of ER, PR, or HER2 in patients with histological diagnosis of breast adenocarcinoma.

In the remainder of this report, we first summarise the results from the model selection and clinical validation study. This is accompanied by a discussion of the results and their implications on the safety and efficacy of the device. Next, we provide insights into the effectiveness of the device across different groups, to identify any biases towards certain subpopulations. This is followed by a distribution analysis of the device predictions. Finally, the Methods section provides a summary of the protocols used to perform the studies, including the description of the experiment design, specifications of the underlying models, details of datasets, performance characteristics used for validation, and statistical considerations.

## Results

### Performance Characteristics Assessment

PANProfiler Breast (ER, PR, HER2) is a software device for assessing the status of ER, PR, and HER2 in breast adenocarcinoma directly from image scan of H&E-stained biopsy/resection slides. The device contains a binary predictive model that returns a negative or positive result if the underlying model is certain about its prediction, or indeterminate if the model is not confident enough (see **Methods: Predictive Thresholds and Confidence Interval**). Typically, the outcome of a binary predictive model is determined by comparing the output score (i.e. probability of a sample being positive) with a predictive threshold, which is generally set to 0.5 (see **Figure 11** of **Methods: Predictive Thresholds and Confidence Interval**). Here, to account for uncertainty and determine a confidence interval of the device, we introduce two predictive thresholds, T1 and T2, and compute the final prediction as follows: if the output score is *greater than or equal* to the *higher* threshold (T2), the device yields a positive prediction. If the output score is *less than or equal* to the *lower* threshold (T1), the device yields a negative prediction. Any output between T1 and T2 is considered indeterminate. This decision mechanism allows a device to predict the status of a case only if it is empirically certain, therefore providing higher safety and efficacy in the clinical setting.

The effectiveness of the device to return confident predictions is measured by “Test Replacement Rate” (TRR), which corresponds to the percentage of predictions performed by the device over all cases. It provides a direct estimation of the number of traditional tests that can be replaced by PANProfiler Breast (ER, PR, HER2) for molecular profiling. Taking the confident predictions into account, we further compute the standard evaluation metrics of accuracy, positive prediction rate (PPV), and negative prediction rate (NPV) (**Supplementary: Detailed Performance Characteristics**). Sensitivity and specificity are not considered in the evaluation, as they are likely to be biased due to the highly unbalanced nature of the data [18]. The impact of this can also be observed in the performance of the state-of-the-art tests used for molecular profiling [19].

### Internal Validation and Model Selection

An internal validation study was conducted to assess the performance of the models to predict ER, PR and HER2 using different model configurations (e.g. different backbones, see **Methods: Pre-processing Pipeline and Training Details**) to determine the final models that would be integrated into the device for the clinical validation. Performance metrics previously mentioned were considered during the evaluation. A total of 8 different configurations were evaluated for each biomarker, repeated three times, each on a randomly sampled subset of the same dataset. 72 models (8 configurations x 3 repeats x 3 biomarkers) in total were trained as part of this internal model selection study. Violin plots showing the distributions of certain metrics computed across all configurations are presented for each biomarker in **Figure 3**. Scatter plots showing the correlation of TRR with accuracy, PPV, and NPV are given in **Figure 4**.

**Figure 3:**
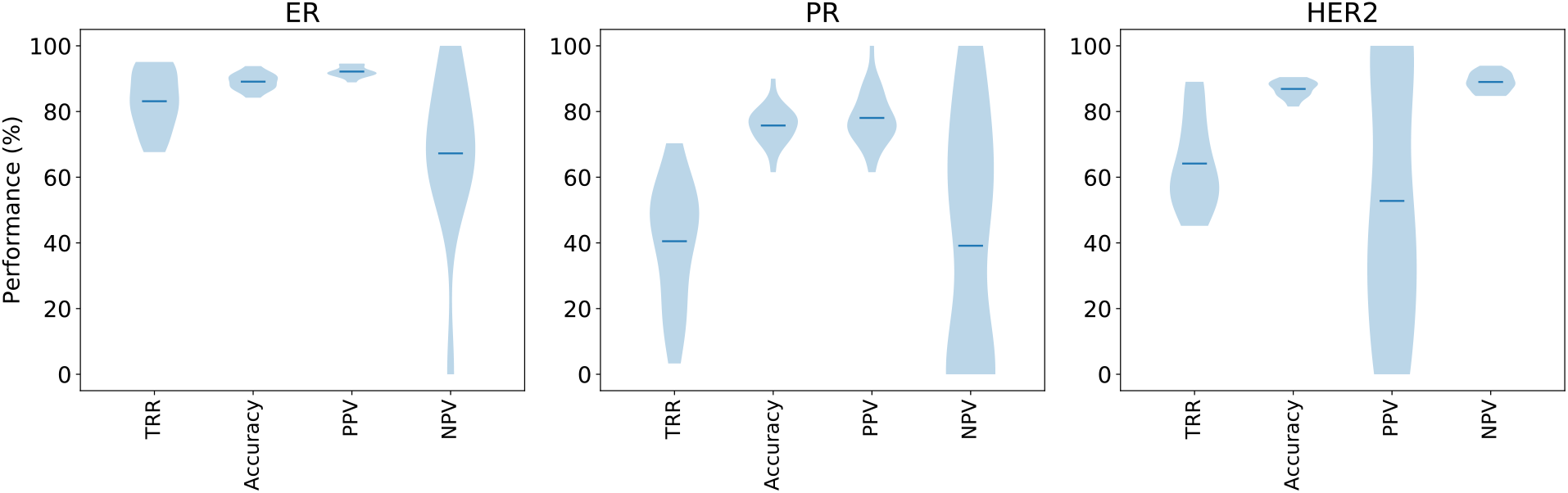
ER, PR, and HER2 violin plots showing the probability density of evaluation metrics acquired from the models trained during the internal model selection study. Each plot shows data from a group of 24 models.

**Figure 4:**
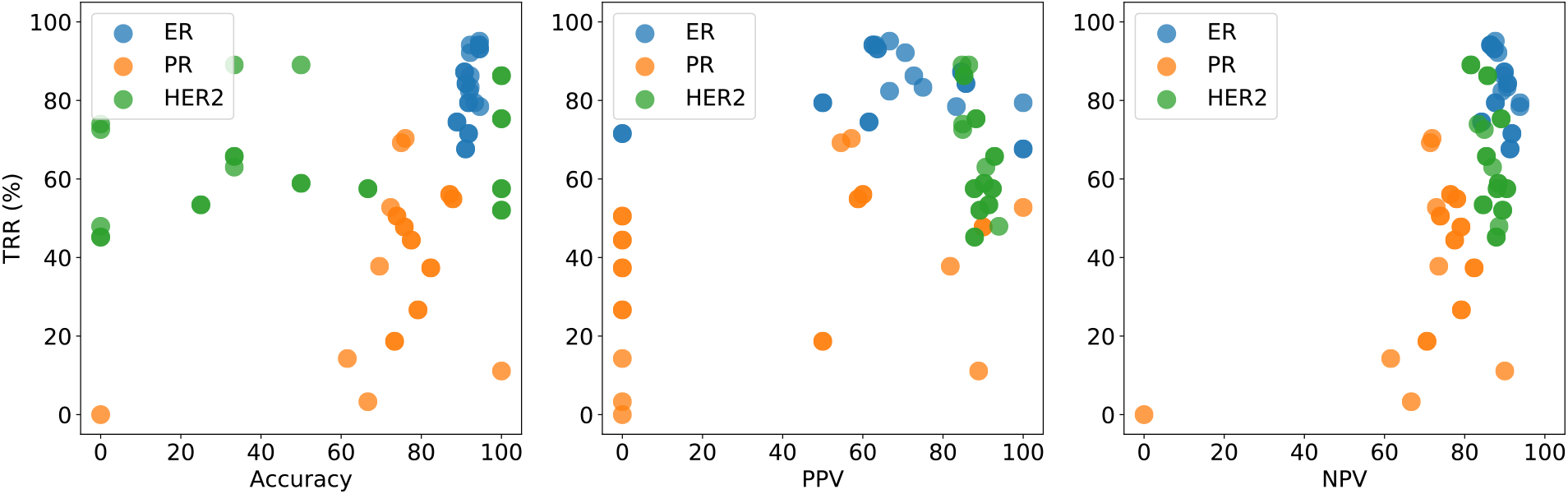
Scatter plots of accuracy, PPV and NPV against Test Replacement Rate (TRR) for ER, PR, HER2 models, validated as part of the analytical performance evaluation study. These performance metrics were simultaneously taken into consideration when selecting the models to be integrated into the device for clinical validation.

Considering the top-performing models for each biomarker, we achieved up to 94%, 80% and 89% accuracy, with a TRR of 78%, 11% and 52% among all the tested cases from an unseen dataset for ER, PR, and HER2, respectively. Detailed performance metrics of the selected models for each biomarker are given in **Table 1**. Metrics for all models included in this study are provided in **Supplementary Table S-1**.

**Table 1:** Performance metrics for the top-performing models in the analytical performance evaluation study. All results are shown as percentages. See **Supplementary: Detailed Performance Characteristics** for the details of validation metrics and refer to **Supplementary Table 1** for the performance metrics of all models included in this study.

| Biomarker | TRR | Accuracy | PPV | NPV |
| --- | --- | --- | --- | --- |
| ER | 78% | 94% | 95% | 83% |
| PR | 11% | 90% | 100% | 88% |
| HER2 | 52% | 89% | 100% | 89% |

The internal validation and model selection study showed the overall performance of the deep-learning models for predicting ER, PR, HER2 directly from routine H&E images within a wide spectrum of metrics. TRR, accuracy, PPV, and NPV on average were mostly stable across different configurations. ER models in particular performed well with TRR, accuracy and PPV roughly between 80 and 95%. A similar trend was followed by PR, but with the expectation of the relatively low performance and high variance of TRR compared to those of the other biomarkers. Despite this, PR models with certain configurations were still very effective and provided viable candidates for the device. For instance, the selected PR model had a low TRR of 11% but performed well with accuracy, NPV and PPV of over 88%. HER2 models showed comparable accuracy to PR with more favourable TRR but relatively low stability for PPV. Considering the safety requirements of the device, we argue that the benefit of including the positive prediction to HER2 is marginal as only a small majority of reliable cases are likely to be predicted as positive, potentially resulting in a low PPV [18]. We, therefore, decided to make HER2 a negative-only predictor, meaning that the device was only allowed to provide negative or indeterminate predictions for HER2 in the clinical performance evaluation study.

### Clinical Performance Evaluation

A clinical performance evaluation study was conducted to assess the effectiveness and safety of the device based on the ER, PR, HER2 models selected in the internal validation study. Models were first calibrated to adjust the predictive thresholds (see **Methods: Model Calibration**) and then evaluated on the unseen test dataset, which included 370, 381, and 296 cases for ER, PR, and HER2 respectively. The results for key performance metrics are provided in **Table 2** for all three biomarkers.

**Table 2:**
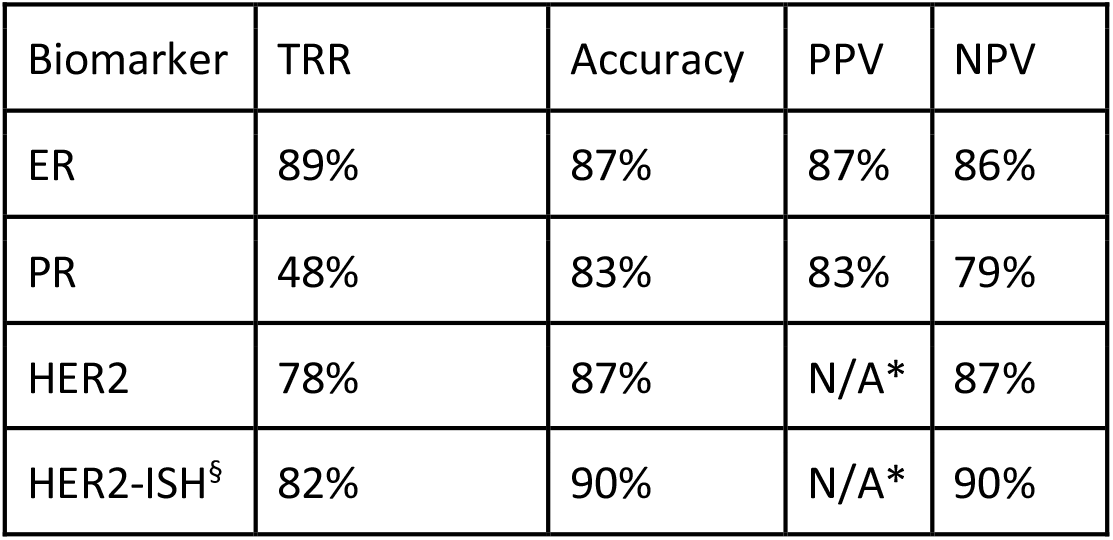
Final performance metrics for each biomarker computed on the test dataset used in Clinical Performance Evaluation Study. *: In the clinical study, the method was only allowed to predict negative results for HER2, therefore PPV is not applicable for analysis. §: HER2 concordance of the device with ISH results.

For ER, the device achieved a good performance simultaneously for every considered metric. PR follows a very comparable trend with regards to accuracy, PPV, and NPV. Despite having a relatively low TRR, its effectiveness is still highly similar to that of ER, with only an overall 4 to 7% decrease in accuracy, NPV and PPV. A high level of TRR was measured for HER2 and its general performance was mostly on par with ER. Since the method was only allowed to predict negative results for HER2, PPV was not applicable for analysis. We also evaluated the concordance of the device to the ISH tests carried out for the HER2 profiling and noticed a slight increase in accuracy, NPV and TRR. Overall, these results show the effectiveness of PANProfiler Breast (ER, PR, HER2) as a predictive tool that can largely eliminate the need for the current standard tests for profiling the molecular biomarkers of breast cancer.

### Confounding Factor Analysis

The clinical validation results in **Table 2** provide an overview of the overall performance of the device on the entire validation population. In this section, we further investigate the effectiveness of the device across different groups, to identify any biases towards certain subpopulations. Towards this end, we have divided the test sets into subgroups based on age, clinically relevant molecular subtypes (HER2-enriched, Luminal A, Luminal B and triple-negative), pathology laboratories (sites), cancer staging (overall), and ethnicity. **Figures 5, 6, and 7** respectively show the performance of ER, PR and HER2 models across these subpopulations.

**Figure 5:**
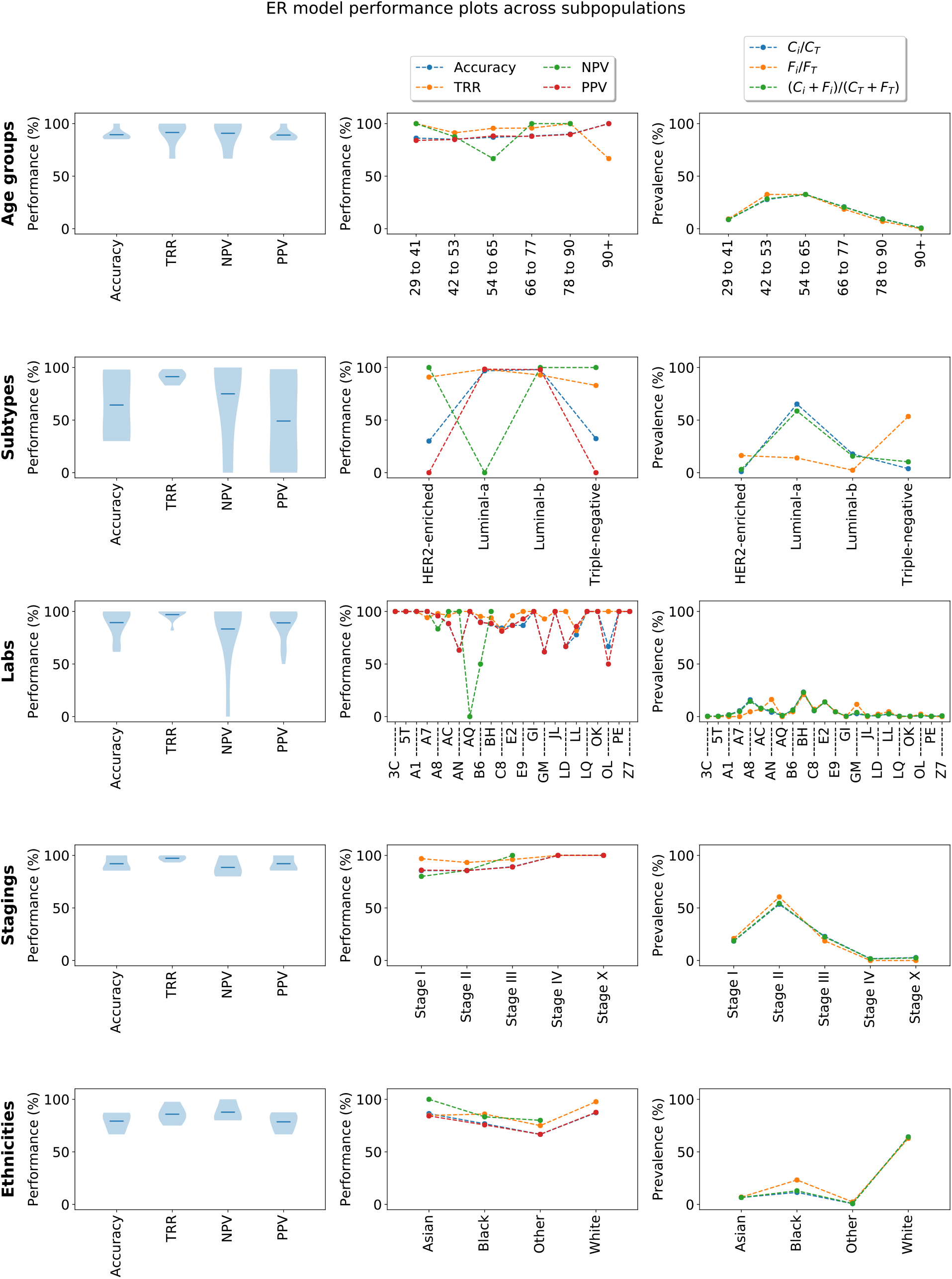
Performance plots of the ER model across subpopulations confounded by (from top to bottom) age, molecular subtype, ethnicity, pathology laboratory (site), cancer stage, and ethnicity. **Column 1:** Violin plots showing the performance distribution across population groupings. Each distribution here has the same sample size as the number of categories in their respective grouping, e.g. for ethnicity, each distribution contains 4 data points, since there are 4 ethnicities in the dataset (Asian, Black, Other, White). **Column 2:** Curves showing the overall performance of the model across different categories. **Column 3**: Prevalence curves of grouping in the populations where the model made 1) correct predictions (blue), 2) false predictions (yellow) or 3) any prediction (green). Here, Ci denotes the number of cases in category i with correct predictions, whereas CT denotes the total number of cases with correct predictions in the whole population. Similarly, Fi denotes the number of cases in category i with false predictions and FT denotes the number of cases with false predictions in the whole population. For curves in columns 2 and 3, it should be noted that except in the case of age groups and staging, the orderings of the categories across the x-axis are completely arbitrary, and the lines between them were added for the sole purpose of visual aid. The laboratory names in row 2 were abbreviated for visual purposes. The full names are provided in **Supplementary Table S-2**. The molecular subtypes of breast cancer were determined based on the following classification --Luminal A: ER+ and/or PR+ and HER2-. Luminal B: ER+ and/or PR are positive and HER2+. Triple-negative (basal-like): ER-, PR-, and HER2-. HER2-enriched: ER+, PR+ and HER2+ [20]. Stages I to IV correspond to standard AJCC pathologic tumour stages. Stage X represents the cases where the status of staging could not be determined.

**Figure 6:**
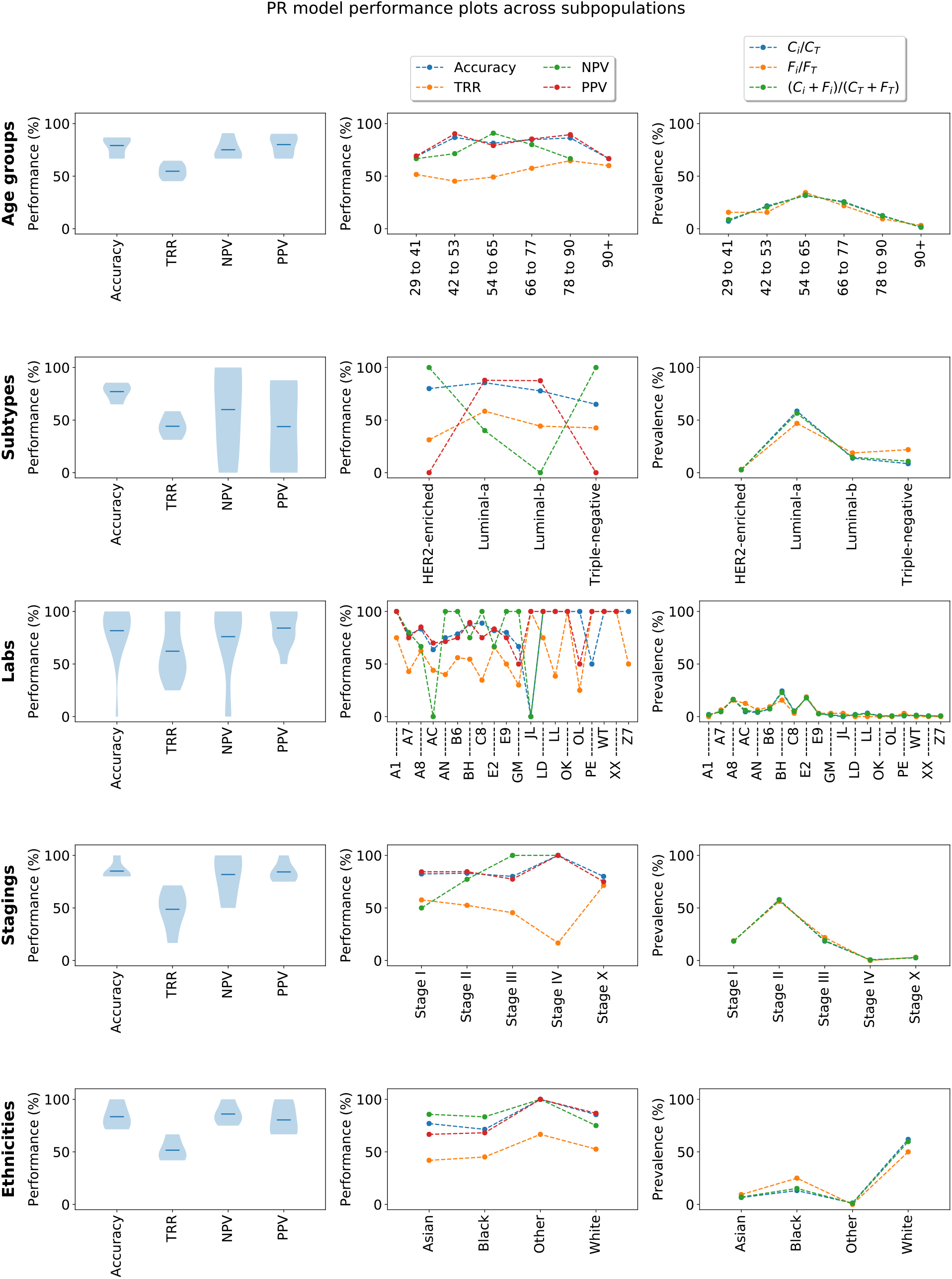
Performance plots of the PR model across subpopulations confounded by (from top to bottom) age, molecular subtypes, ethnicity, pathology laboratories (sites), cancer stage, and ethnicity. Please refer to the caption of **Figure 5** for detailed explanations of the plots.

**Figure 7:**
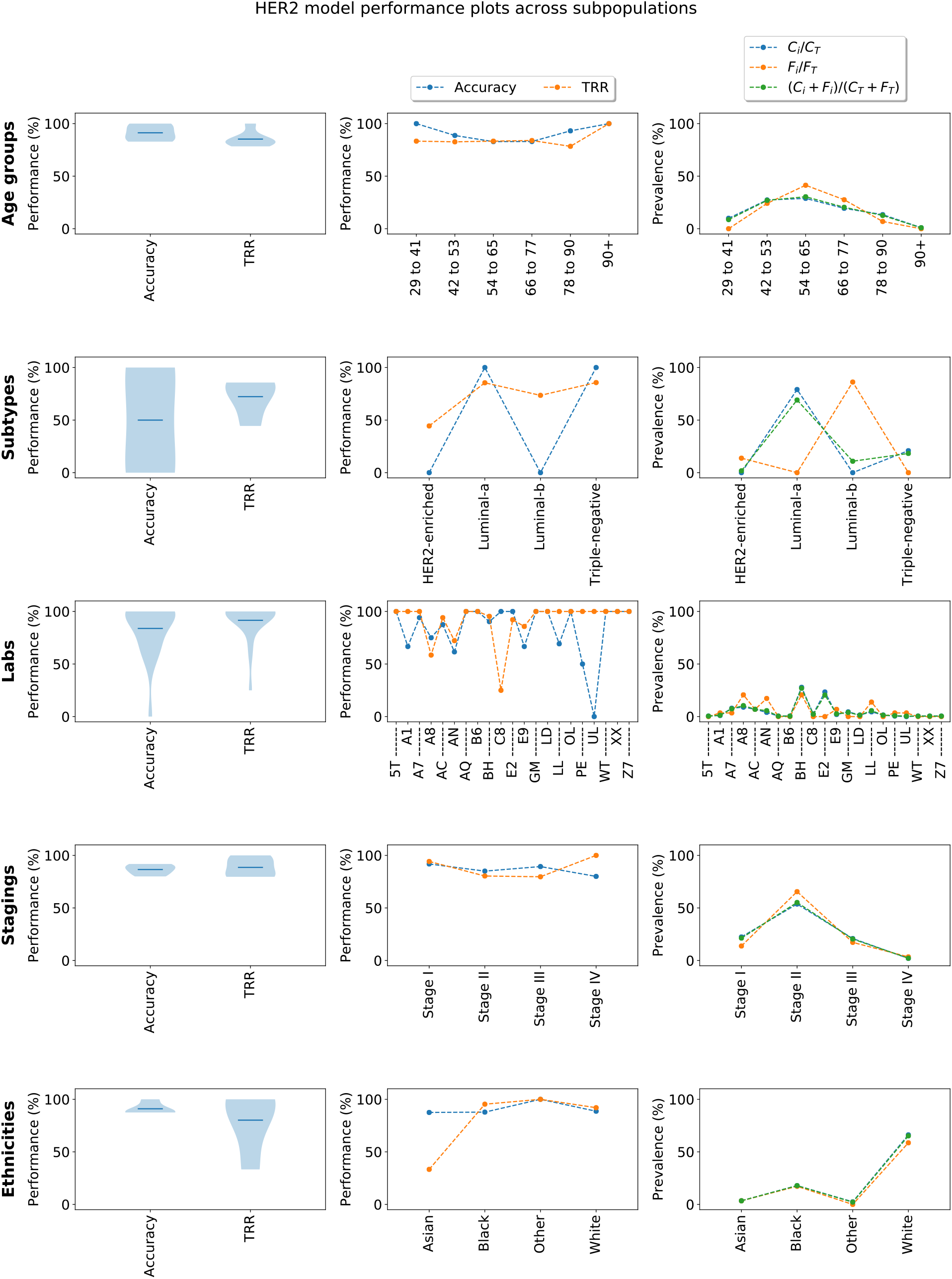
Performance plots of the HER2 model across subpopulations confounded by (from top to bottom) age, molecular subtypes, ethnicity, pathology laboratories (sites), cancer stage, and ethnicity. Please refer to the caption of **Figure 5** for detailed explanations of the plots. Unlike **Figures 5 and 6**, only accuracy and coverage are shown for the HER2 model since the HER2 model can only perform negative predictions.

**Column 1** of **Figure 5** shows the distribution of performances across age groups (**row 1**), molecular subtypes (**row 2**), laboratories (**row 3**), cancer stagings (**row 4**) and ethnicities (**row 5**). In general, the performances are more variable across labs and molecular subtypes but highly consistent across age groups, cancer stages and ethnicities. The variability of the performances across molecular subtypes can be seen in **column 2, row 2**. The inter-laboratory difference in performance can be attributed to the sample size of each laboratory, which tends to have a very high variance. For instance, BH has 77 cases while many others have only a single case, making their performance metrics either 0 or 1. The variation in performances across subtypes can also be attributed to the inherent class imbalance of the category. Triple-negative contains only negative classes across all biomarkers, HER2 enriched ER-, PR- and HER2+, Luminal A ER+ and/or PR+ and HER2-; and Luminal B ER+ and/or ER- and HER2+ [20].

One group worth noting is HER2-Asian, where the TRR seems to have dropped considerably compared to the other ethnicities. Similarly, for the ER-90+ age group and the PR-Stage IV group, the models seem to be more conservative with their predictions, yielding a noticeable drop in TRR. However, this trend can also be attributed to the sizes of these groups (**Supplementary Table S-3**), which tend to be smaller than the other subpopulation partitions, making the performance characteristics less reliable.

The prevalence figures (**column 3**) show the relative rate of correct and false predictions across different groupings of a subpopulation. The plots describe how the correct predictions are distributed across the subpopulations (**blue curve**), how false predictions are distributed across subpopulations (**yellow curve**) and how these compare to the distribution of the predicted population (**green curve**, which is the combination of blue and yellow). In an ideal scenario, we expect the three prevalence curves to be similar across categories (age group, subtype, lab etc), as this indicates that the false predictions are distributed evenly across population types. On the flip side, a deviation between the curves indicates a performance bias. Overall, no significant deviation across subpopulations is observed in all biomarkers, except for subtypes. In this instance, false prediction prevalence for Luminal A and triple-negative in **Figure 5** (ER plots) largely deviates from the overall trend, which might indicate that the ER model tends to return fewer false predictions in Luminal A cases but more often makes false predictions within the triple-negative group. This can be attributed to the ER model being more effective at predicting ER-positive status than negative.

Similar trends can be observed for the prevalence curve of the HER2 model in predicting the markers across molecular subtypes, where it appears to be less common for the model to make false predictions within the Luminal A group. This trend appears to change direction for Luminal B, where the model tends to return false predictions at a higher rate, which can be due to the HER2 model being a better negative predictor. It should also be noted that the number of correct HER2 predictions in Luminal B and HER2-enriched is by definition 0 since we only make negative HER2 predictions and the HER2 status in those subtypes are always positive. Compared to ER and HER2, the PR model appears to be relatively balanced, as we cannot observe much deviation.

### Distribution Analysis of Model Predictions

To better understand the distribution of the device predictions and to see the impact of indeterminate decisions on the device performance, we plotted the histograms of prediction scores for the ER, PR, and HER2 models. Prediction score roughly measures the model confidence of predicting a case as a positive or negative (see **Figure 11** of **Methods: Predictive Thresholds and Confidence Interval)**. The negative and positive predictions were grouped by their corresponding ground-truth labels, acquired from the well-characterised results of the IHC and ISH tests (**Figure 8**). The plots were further subdivided into three zones, representing negative, indeterminate and positive predictions, where the indeterminate zone is defined by the low and high predictive thresholds.

**Figure 8:**
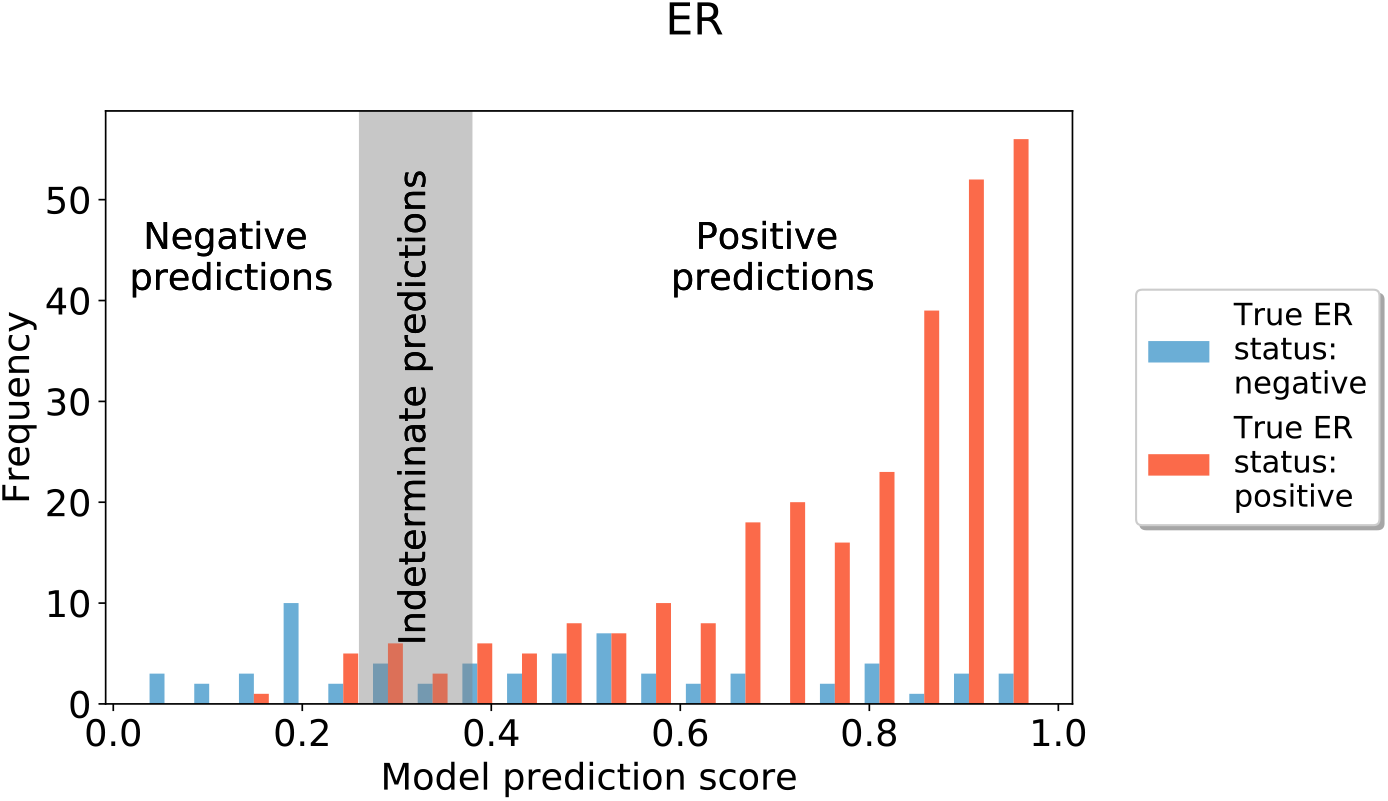
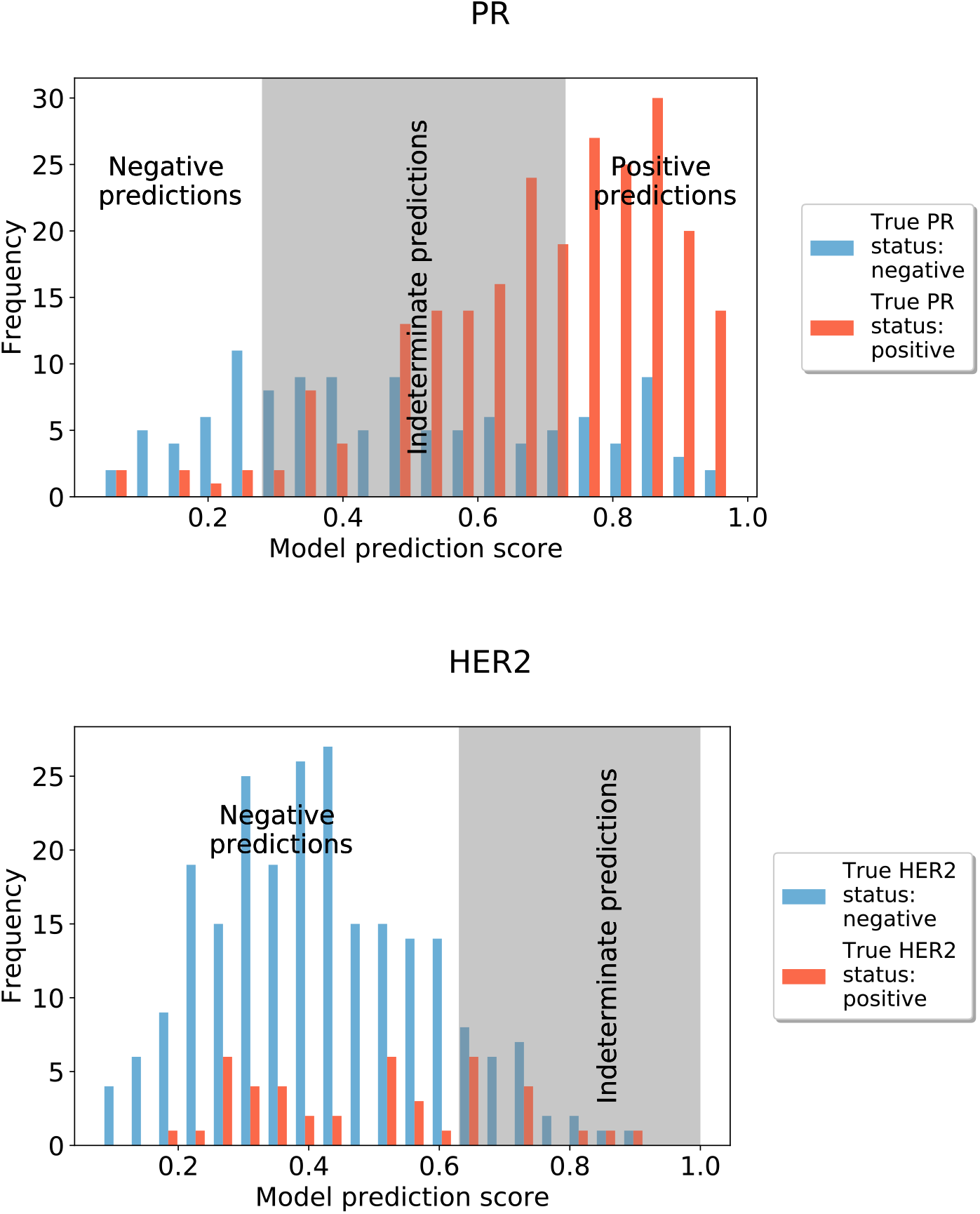
Histograms of model prediction scores for ER, PR, and HER2. Blue and red bars represent the ground-truth labels, acquired from the well-characterised results of the IHC and ISH tests. A grey zone is bound by the predictive thresholds and corresponds to the indeterminate zone. Based on its relative location to the grey area and its true value, a prediction can fall into one of the four categories of true positive (**TP**), true negative (**TN**), false positive (**FP**), or false negative (**FN**). **TN**: Blue bars to the left of the grey area. **FN**: Red bars to the left of the grey area. **FP**: Blue bars to the right of the grey area. **TP**: Red bars to the right of the grey area. Blue and red bars within the grey area represent cases where the model does not make a prediction. Since HER2 is only allowed to perform negative predictions, no positive predictions are shown. ER prediction model performs very well in separating negative and positive classes. While the PR prediction model does not achieve as well separation between positive and negative classes as PR prediction’s, the overall separation between the positive and negative samples can still be seen in the histogram. Given that the HER2 model could not well separate negative and positive classes, only negative classes are being called by the model.

The distributions of true positives (TP), false positives (FP), true negatives (TN), and false negatives (FN) of ER, PR and HER2 predictions reflect the overall results presented in **Table 2**. Reflecting the results on the table, a very low FN and FP rate for ER/PR and HER2 are observed from the distribution. Logically, there is an inverse relationship between TRR and the width of the indeterminate zone: the more confident a model gets with its predictions, the higher TRR it yields, i.e. the smaller the intermediate zone is. This serves as a safety mechanism for the device, as rather than performing an uncertain prediction that is likely to be a false positive or negative, it simply decides to return no outcome, allowing the receptor status to be determined with the standard of care procedures. The histograms (**Figure 8**) also clearly show that the positive and negative prediction distributions are highly aligned with the prevalence of the underlying data, which is typically highly unbalanced, e.g. according to the 2019 report by the Royal College of Pathologist [18], the population-based positive class ratios for ER, PR and HER2 status are 4.24, 2.12, and 0.17, respectively. For instance, the ER and PR models tend to predict more positives and the one for HER2 is likely to predict more negatives.

### Interpreting Model Predictions

To better understand how deep-learning models determine the molecular status from H&E images and to obtain some insights towards identifying certain morphological features that can be associated with molecular alterations, we plotted the highest and lowest scoring tiles (i.e. most confident positive and negative predictions) for a selected set of 12 whole slide images (WSI) from each of the ER, PR and HER2 test sets used in the clinical validation study. The visualisations for the ER model are provided in **Figure 9**. We show three examples for each of true positive (TP), false positive (FP), false negative (FN) and true negative (TN). Similar visualisations are created for the PR and HER2 models and are shown in **Supplementary Figures S-1** and **S-2**, respectively.

**Figure 9:**
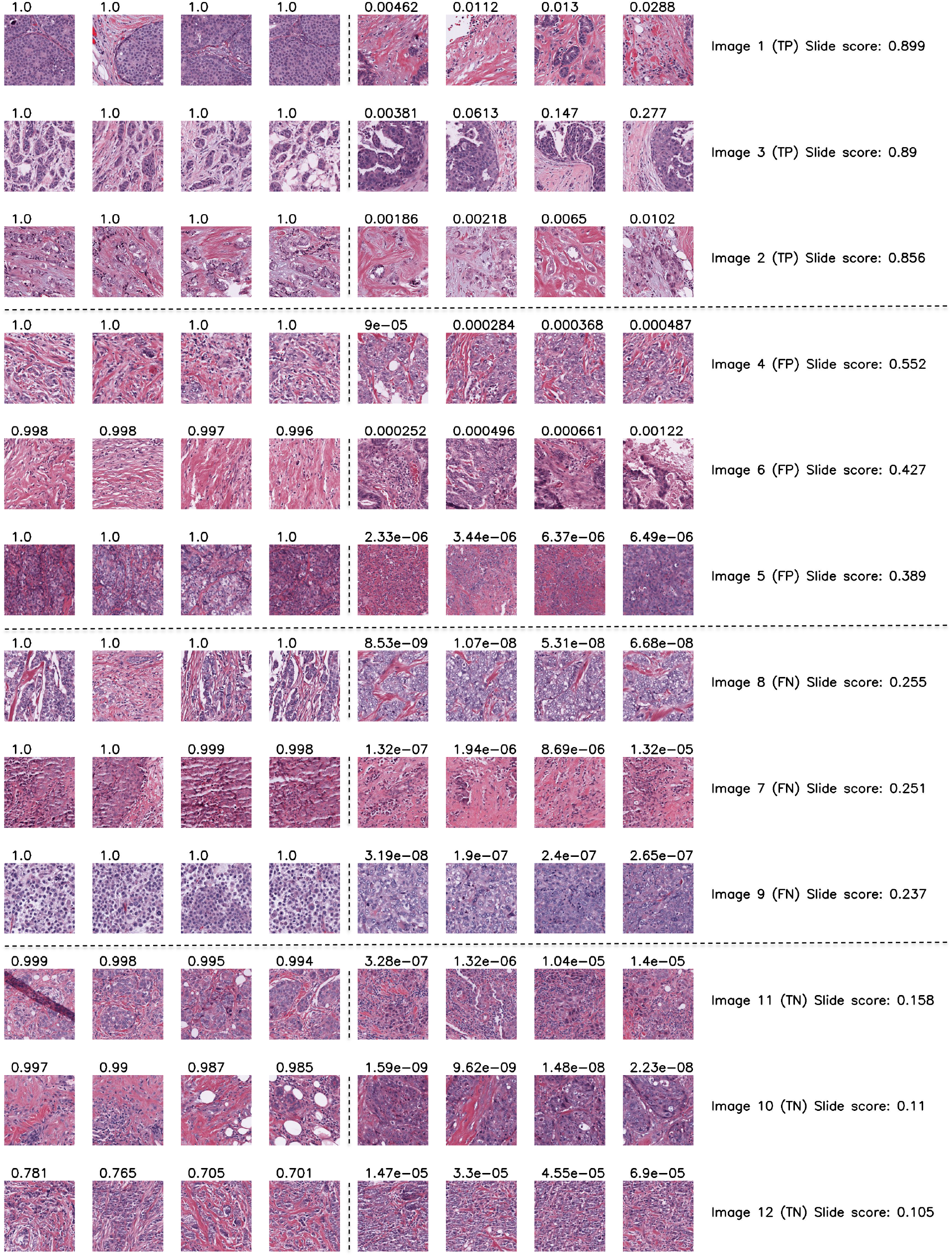
Highest and lowest ranking tiles acquired from a selected set of 12 whole slide images (WSI) from the ER test set. Each row corresponds to a WSI that is classified as one of the following categories based on the model prediction and the ground-truth receptor status: true positive (TP), false positive (FP), false negative (FN), and true negative (TN). The leftmost four tiles in each row are the ones with the highest score whereas the rightmost four tiles represent those with the lowest score. The values above the tiles correspond to the tile-level model output while the values alongside the image name correspond to the slide scores, which are used by the device to return the final prediction.

In Figure 9, one can see the distinct patterns between the highest and the lowest ranking tiles (first half and second half in each row, respectively), indicating that the model can capture certain visual features from the images when distinguishing a positive or negative tile. Ideally, for the correctly predicted cases (i.e. TP and TN), the majority of the slide scores should be close to 1 for positives and close to 0 for negatives, whereas, slide scores should be closer to the indeterminate zone when the model is performing wrong predictions (i.e. FP or FN cases). However, this trend is not strongly observable in our figures, especially when interpreted together with the distributions of the model predictions (Figure 4). One can note that the ER model is the closest to the ideal case, where the histogram of its positive predictions shows a left-skewed distribution (i.e. most predictions are close to 1).

Another key observation from Figure 9 is the morphological variability across the high-ranking tiles of different slides. We are now in the process of inspecting these tiles to identify any associations between the tumour morphology and the model predictions. Future work will also explore the internal representations of the model to reveal any histopathological patterns linked to molecular alterations.

## Methods

### Pre-processing Pipeline and Training Details

A proprietary convolutional neural network (CNN) model was used to determine the status of molecular profiles for breast cancer. The workflow of the prediction of breast cancer molecular profiles from H&E images with deep learning is illustrated in **Figure 10**. Each model was trained on a set of 256×256 tiles acquired from whole slide images (WSI) stained with H&E. In the first step of the pipeline, the image is broken down into tiles. A standard deviation filter is used to eliminate the background tiles which do not contain any relevant information. Then, an independent tumour segmentation model is used to detect the tumour regions and discard non-tumour regions. A WSI was discarded from analysis if it contained fewer than 10 tiles after the filtering process. Macenko colour and brightness normalization [21] was applied to the remaining tiles before they were assigned with a ground-truth label, acquired from the well-characterised results of the IHC and ISH tests. Finally, the pre-processed tiles are used as inputs to train a deep learning model, which learns to make a biomarker prediction.

**Figure 10:**
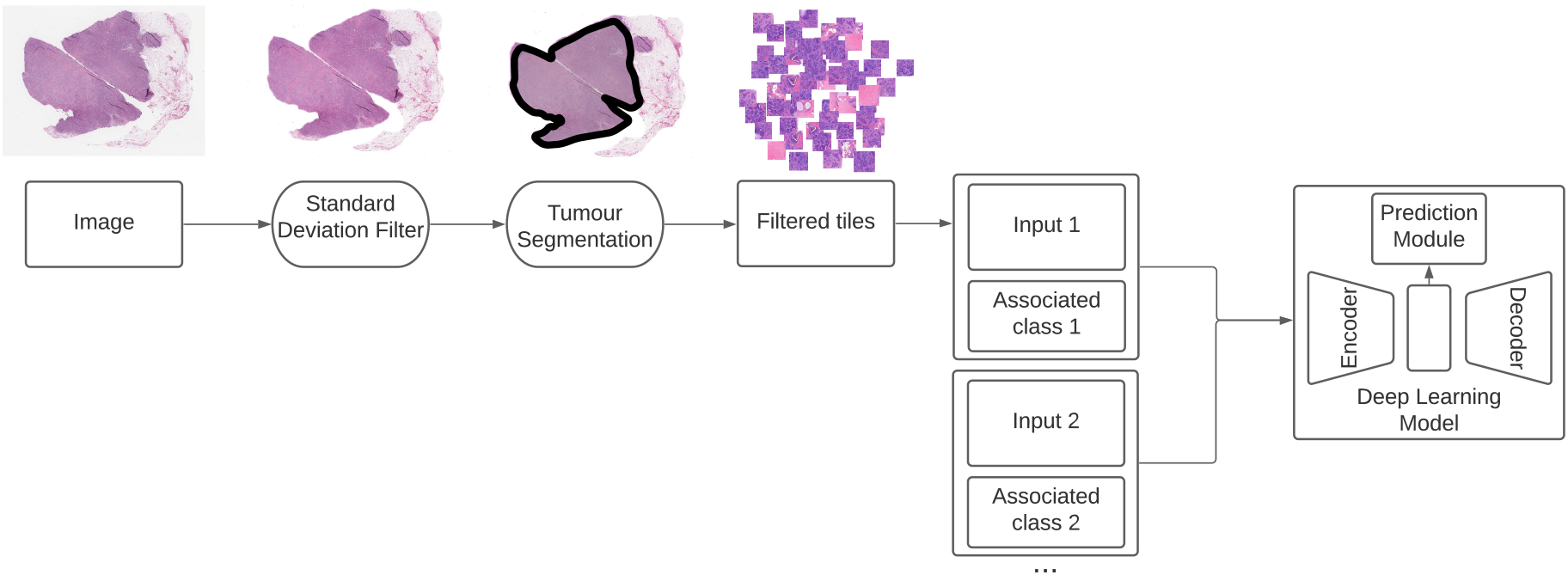
Visualisation of the training pipeline. The prediction pipeline works in a similar fashion in which standard deviation filter and tumour segmentation are used prior to model prediction.

A different CNN was trained for each biomarker, i.e. the final device was formed of three CNNs, each consisting of an encoder extracting features from an input, a decoder and a classification module (see **Figure 10**). The feature extractor can capture the tissue properties within tiles throughout a set of convolutional filters applied to tiles at various layers of depth, effectively encoding the high-level visual features into a d-dimensional feature vector, where d depends on the architecture of the CNN. These vectors are regarded as the fingerprints of the tiles and are submitted to both the decoder and the classification module.

The decoder module takes a *d*-dimensional embedding as input and returns an output of the same shape as the original tile that the embedding represents. It consists of a series of transposed convolutional and upsampling layers, which resembles an inverted copy of the CNN used for feature extraction, known as the encoder. Its purpose is to reconstruct the original tile from the latent vector to achieve better representations of each tile that do not contain irrelevant features.

In parallel to the decoder, the feature vector representative of each tile is also submitted to the classification module, which consists of a fully connected layer. The output classification score is then compared to the label of the tile’s parent WSI. All modules are trained end-to-end and each tile is given a score, i.e. confidence of it being positive. Finally, the scores of all the sampled tiles from each WSI are aggregated via mean pooling to produce the final slide-level scores.

### Target Patient Groups

The following target patient groups of both sexes were included in the validation studies: diagnosed cases of invasive breast carcinomas including bilateral carcinomas, histologically distinct ipsilateral carcinomas or widely separated carcinomas considered to be separate synchronous primary tumours, recurrent breast carcinoma, metastasis of breast carcinomas, ductal carcinoma in situ.

### Datasets

Both validation studies were conducted on digitally scanned images of formalin-fixed paraffin-embedded core breast biopsies and resection sections. Data was acquired from 1) the Cancer Genome Atlas (TCGA) open-access dataset for the breast adenocarcinoma (BRCA) study [22] (i.e TCGA-BRCA) and 2) a proprietary dataset by a private clinical data provider (e.g. BioIVT). All samples were following the inclusion criteria described in Target Patient Groups.

Prior to going to the internal validation phase, the model was pre-trained on a wider internal dataset. For the internal validation and model selection study, we used the entire dataset of BioIVT and a subset of TCGA-BRCA. To ensure confounder independence and show generalisability during validation, different TCGA participant labs were chosen for model selection and clinical validation. We selected the labs such that the resulting datasets reflect the population distribution of population groupings including molecular subtypes. The details of the analytical evaluation datasets, including the number of positive and negative cases for each biomarker, are given in Table 4.

**Table 4:** Details of the dataset used for the analytical validation study. Please note that the number of cases differ for each biomarker as the actual receptor status may not always be available for all cases/biomarkers. Values in parenthesis represent the number of positive/negative cases.

| Dataset | ER (+/-) | PR (+/-) | HER2 (+/-) |
| --- | --- | --- | --- |
| TCGA-BRCA | 342 (272/70) | 339 (232/107) | 328 (49/279) |
| BioIVT | 435 (343/92) | 360 (228/132) | 269 (91/178) |
| Total | 777 (615/162) | 699 (460/239) | 597 (140/457) |

For the clinical evaluation study, we used a subset of the TCGA-BRCA dataset which only contained images from the laboratories that were not included in the model selection study. The details of the dataset, including the number of positive and negative cases for each biomarker, are given in Table 5.

**Table 5:** Details of the dataset used for the clinical validation study. Please note that the number of cases differ for each biomarker as the actual receptor status may not always be available for all cases/biomarkers. Values in parentheses represent the number of positive/negative cases.

| Dataset | ER (+/-) | PR (+/-) | HER2 (+/-) |
| --- | --- | --- | --- |
| Total | 648 (503/145) | 648 (437/211) | 560 (134/426) |

### Predictive Thresholds and Confidence Interval

The outcome of a binary model is determined by comparing the output score (i.e. confidence of a sample being positive) to a predictive threshold, yielding either a positive or negative result. PANProfiler Breast (ER, PR, HER2), on the other hand, can also return an indeterminate result if the prediction of the device is not within a “confidence interval”. To determine this confidence interval, we define two predictive thresholds, T1 and T2 (where T1 < T2), as opposed to a single threshold used in standard predictive models. The definition of T1 and T2 and their impact on the performance of a model is illustrated in **Figure 11**.

**Figure 11:**
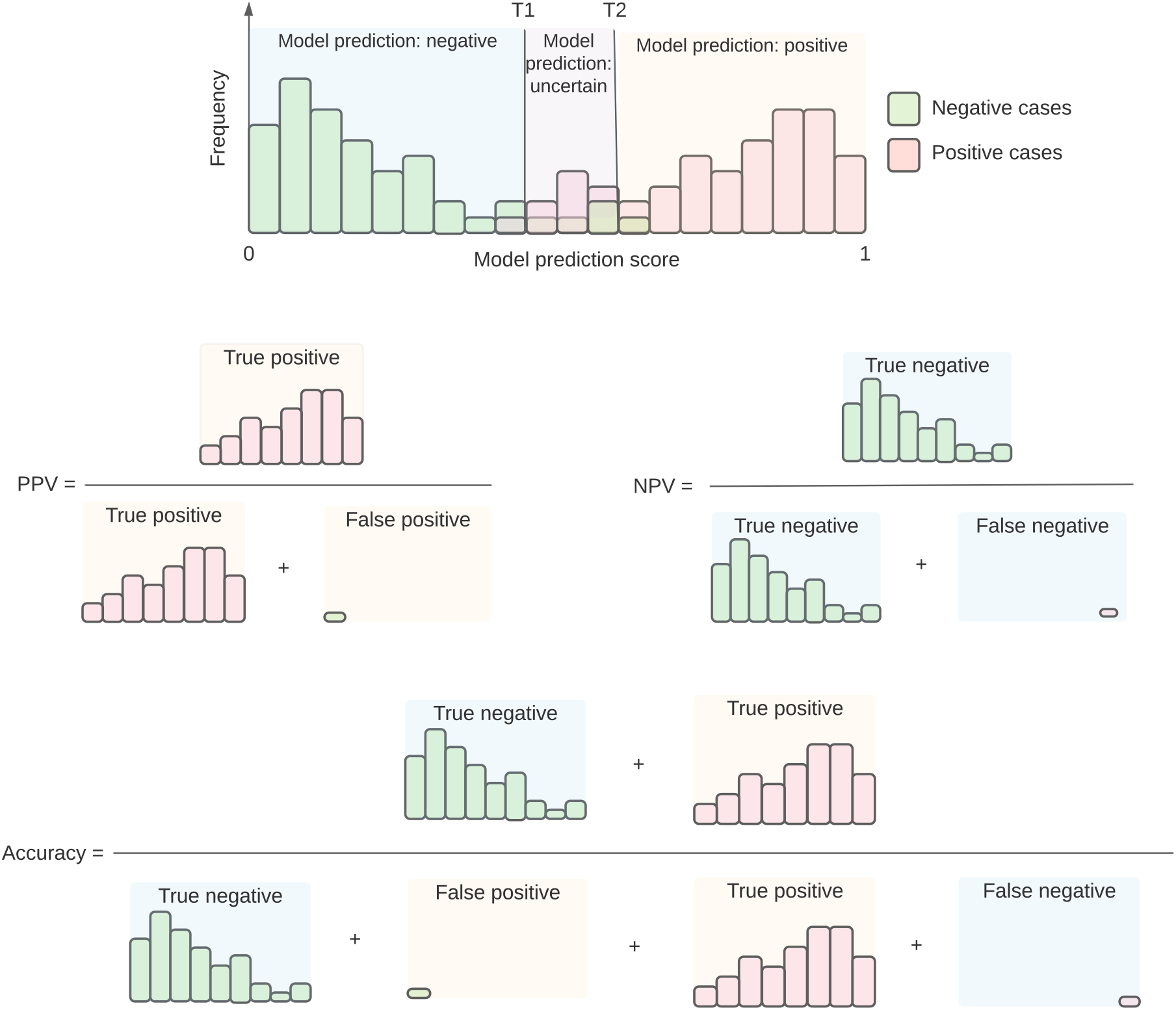
Illustration of the impact of positive and negative predictive thresholds on the sensitivity and specificity of a device.

The predictive thresholds enable the interpretation of the device output as a confidence score. If an output score is greater than or equal to T2 a positive prediction is returned whereas an output with a confidence score less than or equal to T1 corresponds to a negative prediction. Any output between T1 and T2 is considered indeterminate. Ideally, T1 and T2 can be set to the values which yield the highest accuracy for the device. However, in practice, this tends to make the device highly conservative and increases the likelihood of returning indeterminate results, yielding a low TRR. For instance, considering the histogram in **Figure 11**, it is possible to achieve a 100% accuracy if T1 is shifted further to the left while T2 is moved to the right. On the other hand, this comes with increased uncertainty, i.e., a wider grey/intermediate zone. This renders the device unusable in day to day practice. A balance was struck which minimised risk to an acceptable level while allowing a significant number of cases to be predicted reliably. As a result, it is possible to have a device with a high TRR and an accuracy that is acceptable within the safety, efficacy and risk-benefit requirements.

To demonstrate the relationship between the accuracy and TRR in accordance with the predictive thresholds, we conducted an empirical analysis on the models used in the validation studies. The results of the analysis are shown in **Figure 12** with further details provided in **Supplementary: Accuracy-Test Replacement Trade-off**. The predictive thresholds are fitted on a validation set to maximise the TRR whilst maintaining a clinical stakeholder-defined level of accuracy. In **Figure 12**, the blue points represent a set of predictive thresholds and the resulting accuracy-TRR pairs computed during validation. The orange points, on the other hand, correspond to the accuracy and TRR values when the predictive thresholds are applied to an independent test set. It is important to note that, during clinical validation, only the pair of thresholds that maximised the TRR based on a target accuracy of 90% on the validation set was fitted on the final independent test set, which yielded the reported metrics in **Table 2**. Here, we just plot the other predictive thresholds for visualisation purposes.

**Figure 12:**
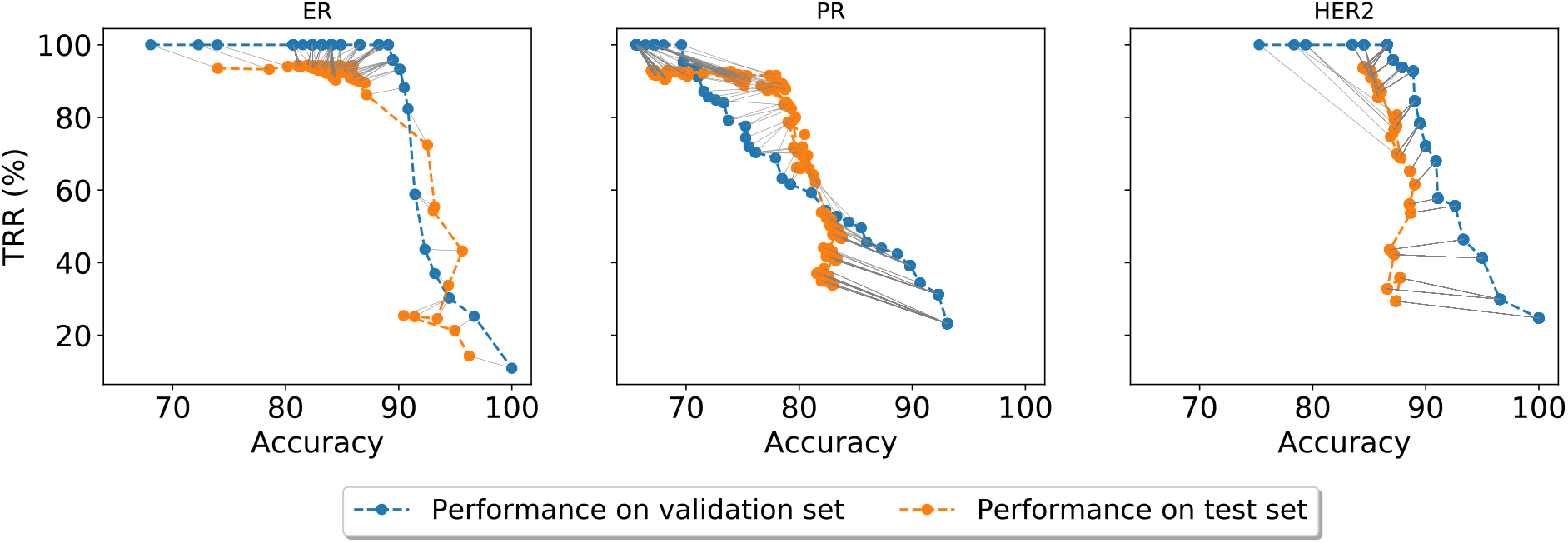
Illustration of the relationship between accuracy and TRR in accordance with the predictive thresholds computed on a validation set (blue line) and applied to a test set (orange line).

As expected, there is an inverse relationship between TRR and accuracy in the validation set. This trend is diminishing on the lower end of the accuracy as decreasing target accuracy does not necessarily increase TRR. This is expected given the observation seen in **Figure 8** where the distribution of true positive and negative samples’ prediction scores are separable. The opposite can also be said about the lower end of TRR: decreasing TRR to achieve more accuracy would have a diminishing return in the lower end of TRR. This is again attributable to the observation seen in **Figure 8**.

The inverse relationship is somewhat maintained moving to the test set. The shift in the trade-off curve is attributable to the shift in population moving from validation to test set. This is mitigated by doing calibration (see **Methods: Model Calibration**).

### Model Calibration

Calibration is a process to ensure that the model safety and efficacy remain on an acceptable level before applying it to a new dataset and involves the following steps: 1) fine-tuning the model and 2) re-computing the predictive thresholds T1 and T2. These thresholds are typically derived during model training to maximise the number of confidently predicted cases within a predefined target accuracy. However, due to potential variations between the source and target domains, a performance drop is expected when the model is applied to a new dataset. This necessitates the calibration of the device, which is typically performed on part of the target dataset (40% in this study). Considering the histogram in **Figure 11**, this process can be viewed as shifting the thresholds T1 and T2 in a way that would maximise the device accuracy while maintaining an acceptable TRR. Here it is worth noting that, once calibration is done, the images used in this process are discarded and the device is evaluated only on the remaining images of the target dataset.

### Experimental Details

#### Model selection study

Multiple pre-trained neural networks were trained for each biomarker using pre-assessed, well-characterised biopsy slide images from the BioIVT and TCGA datasets. 70% of the dataset was used for model training while half of the remaining images (15% of data) was used to determine the best model during training (i.e. internal validation) and the other half (15% of data) was set aside for testing the model on unseen data. It should be noted that this dataset is distinct from the clinical validation data. The sampling algorithm used for selecting cases for each subset was performed at the case level and was constrained to ensure that the population-based distributions of receptor profiles were mostly preserved in the validation and test set. Once these subsets were adjusted to reflect the real-world prevalence of hormone receptors and HER2, the remaining cases were moved to the training set. Each case was represented with only a single WSI in the test set to ensure that the final predictions based on the confidence interval can be computed correctly.

72 models in total were trained as part of this internal model selection study. We considered a total of 8 different configurations for each biomarker, where a configuration was repeated on three randomly sampled subsets of the same dataset (i.e. 24 models per biomarker). Our preliminary results on a hyper-parameter analysis revealed three important parameters that would directly have an impact on the model performance: backbone, CNN-type and validation target metric. Backbone corresponds to the CNN used as a feature extractor and was either “resnet34” [23] or “densenet121” [24]. CNN-type represents the architecture of the underlying CNN, which could be either of “tile-ae”, i.e. tile-level autoencoder or “tile-vae”, i.e. tile-level variational autoencoder. The validation target metric was used to select the best model during training and was set to either AUC or TRR.

After each training epoch, the performance of the target metric on the validation set was monitored and the model with the best validation performance was further evaluated on the test set. The top model for each of the biomarkers was selected based on a combined criterion of accuracy, TRR, PPV and NPV assessed on an independent test set. See **Table 2 of Results** for the performance metrics of the selected models and **Supplementary Table 1** for all 72 models considered for internal validation.

#### Clinical validation study

Cases with a well-characterised IHC receptor status were used to construct biomarker-specific datasets of sizes 648, 648, and 560, for ER, PR, and HER2 respectively. 40% of the samples were used for model calibration while the remaining images (constituting 60% of the study samples) were used for testing the device performance. The calibration set was roughly split into two subsets and one half (i.e. fine-tuning set) was used for fine-tuning the model and the other half (validation set) was used for determining the confidence interval. Similar to the model selection study, a constrained sampling algorithm was used to ensure that population-based distributions of receptor profiles were preserved in the validation and test set. Once these subsets were adjusted to reflect the real-world prevalence of hormone receptors and HER2, the remaining images were moved to the fine-tuning set. The number of images used in the clinical validation study for calibration and test are provided in **Table 6**.

**Table 6:** The details of the calibration and test set used in the clinical validation study. Please note that the number of cases differ for each biomarker as the actual receptor status may not always be available for all cases/biomarkers. Values in parentheses represent the number of positive/negative cases.

| Dataset | ER (+/-) | PR (+/-) | HER2 (+/-) |
| --- | --- | --- | --- |
| Calibration | 278 (204/74) | 267 (178/89) | 264 (89/175) |
| Test Set | 370 (299/71) | 381 (259/122) | 296 (45/251) |
| Total | 648 (503/145) | 648 (437/211) | 560 (134/426) |

During calibration, after fine-tuning the model, T1 and T2 were set to the values that maximised TRR, while keeping the accuracy computed on the validation set at 90%. Once the confidence interval was determined, the final clinical performance of the device was evaluated based on the independent test set.

### Device clinical workflow

For clinical performance characteristics, the final selected model was deployed onto the PANProfiler Breast (ER, PR, HER2) device and validated against clinical performance characteristics (some of them described in **Supplementary: Detailed Performance Characteristics**). The device consists of a programmatic endpoint (application programming interface, API) that provides a connection to pathology lab information and picture archiving and communication systems (LIM/PACS). Additional usability studies were conducted to validate the device usability among board-certified pathologists.

### Statistical Considerations

To assess the statistical significance of the sample size used in the clinical validation study, we performed a minimum sample size estimation approach [25] that takes into account the proportion of positive samples (defined as *X/n*, where *X* is the number of positive samples and n denotes the size of a dataset), a margin of error (set to 5%) and a standard confidence interval (set to 95% CI based on a z-score of 1.96). For the whole clinical validation dataset (**Table 6**) the minimum sample size per biomarker was determined as 264, 340 and 280 for ER, PR, and HER2, respectively. Considering the independent test set only, the minimum sample size per biomarker was determined to be 236, 334, 196 for ER, PR and HER2, respectively.

## Conclusions

This paper provided the performance evaluation results of PANProfiler Breast (ER, PR, HER2), a medical predictive device that was developed for the molecular profiling of ER, PR, and HER2. The status of these biomarkers is used to determine the prognosis of breast cancer and to select the most suitable therapy for a patient. IHC and ISH, the conventional tests used for molecular profiling of ER, PR, and HER2, pose a bottleneck in the current standard practice, as they account for a large proportion of the high turnaround time for breast cancer biopsy diagnosis due to additional work and pathologist time required in IHC/ISH preparation and analysis. PANProfiler Breast (ER, PR, HER2), on the other hand, is built upon deep-learning models that can be used to infer the status of ER, PR and HER2 directly from digitally scanned H&E-stained biopsy/resection slides. Results published in this paper show the effectiveness of the device as a predictive tool that can largely eliminate the need for the current standard tests for profiling the molecular biomarkers of breast cancer.

## Acknowledgement

The results shown in this study are partially based upon data generated by the TCGA Research Network: https://www.cancer.gov/tcga.

The authors would like to acknowledge Oscar Maiques Carlos who has been reviewing the visual elements of the experiment as part of this project.

## Authors contribution

P.R-L. conceptualised and led the research. P.R-L. and D.M. designed the study experiment. S.A., X.L., J.H., C.B. and P.R-L. developed the core deep learning and analytical methods used in the experiments and the medical device. J.S., J.H., A.G. and X.L. developed the medical device. K.B., A.M., D.M., A.G., J.H., J.S., C.B. and T.D. verified and validated the medical device. S.A. and J.S. conducted the validation experiments for the manuscript. K.B., A.M., A.G., S.A., J.S., J.H., X.L., C.B. documented verification, validation and experiment protocols and reports. P.R-L., S.A. and D.M. validated experiment results. S.A., X.L. and P.R-L. analysed experiment results. S.A., P.R-L. and X.L. wrote the manuscript. Everyone discussed the results and reviewed the manuscript.

## Supplementary

### Understanding Model Predictions for PR and HER2

**Figure S-1:**
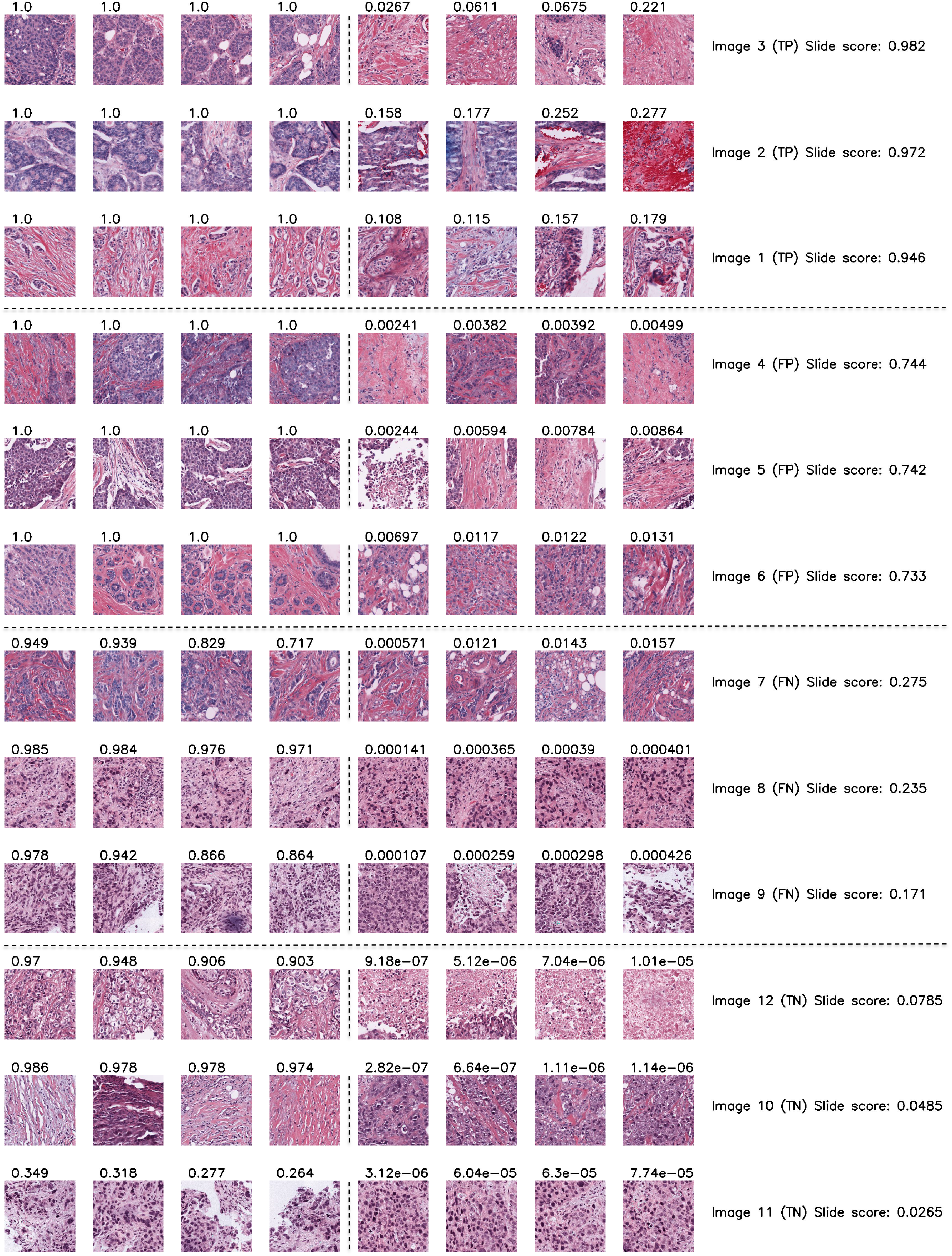
Highest and lowest scoring tiles acquired from a selected set of 12 whole slide images (WSI) from the PR test set. Please refer to the caption of **Figure 9** for a detailed explanation of the visualisation.

**Figure S-2:**
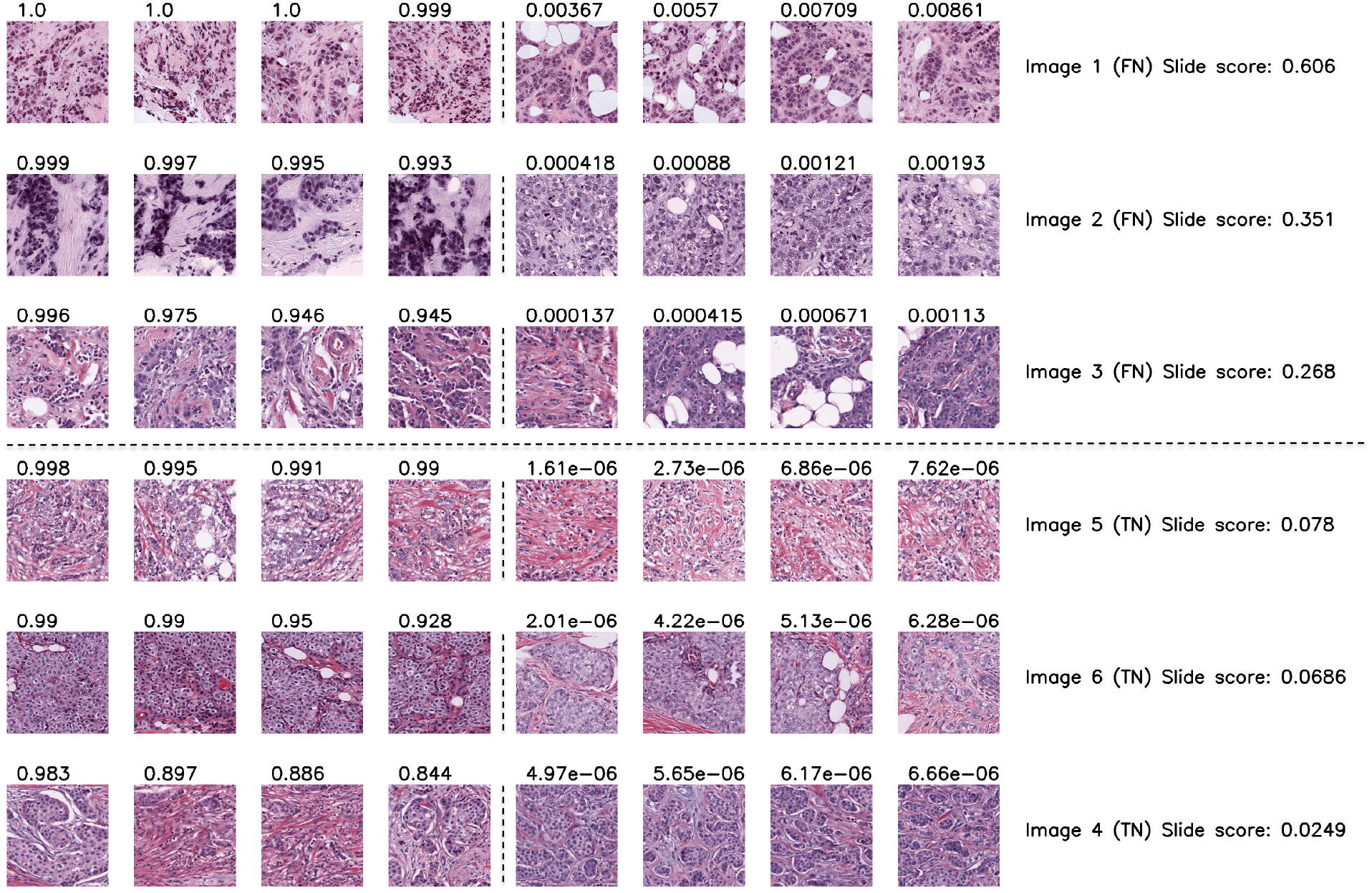
Highest and lowest ranking tiles acquired from a selected set of 6 whole slide images (WSI) from the HER2 test set. Please refer to the caption of **Figure 9** for a detailed explanation of the visualisation. Unlike **Figure 9** and **Figure S-1**, only True Negative (TN) and False Negative (FN) cases are shown, since the HER2 model can only perform negative predictions.

### Accuracy-Test Replacement Trade-off

We conducted a post-validation analysis to demonstrate the relationship between accuracy and TRR in accordance with the predictive thresholds. The results of the analysis are shown in **Figure 11**. In this section, we provide the details of how the plots in **Figure 11** were generated.

During training or calibration, a predictive model M is deployed onto a validation dataset S that is specifically used for computing the predictive thresholds. This dataset only contains images that have not been seen by the predictive model M. For each image, M produces a confidence score between 0 and 1 and given any pair of threshold values (a, b) ⊆[0, 1] we can compute a set of predictions with confidence on the dataset S, considering a prediction to be uncertain if its score lies within (a, b), that is

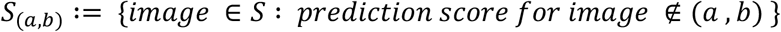

We can then define accuracy (*acc*) and TRR (*trr*) as follows:

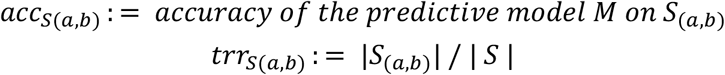

Furthermore, since |S| is finite, the set of “achievable” accuracy values

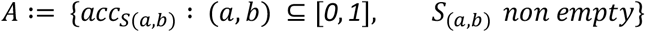

also has a finite number of elements, so for any value of accuracy *α*in *A* there exist thresholds (a(*α*), b(*α*)) which optimise for TRR, such that

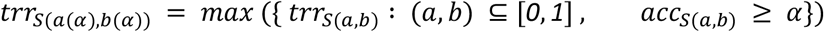

This allows us to plot the relationship between accuracy and test-replacement rate during the threshold fitting process on the validation set *S*, as shown by the blue lines in Figure 10. Here the x-coordinate of each point on the line is an “achievable” value of accuracy for *S*(*a*(*α*), *b*(*α*))and the corresponding y-coordinate is its optimised TRR. We then apply all pairs of predictive thresholds (*a*(*α*), *b*(*α*)) to *T* and plot the resultant accuracy and test replacement values (*accuracy*_*T*(*a*(*α*),*T*(*α*))_, *trr*_*T*(*a*(*α*),*T*(*α*))_) as points on the orange line. The grey segments demonstrate this application of thresholds by connecting points with coordinates (*acc*_*T*(*a*(*α*),*T*(*α*))_, *trr*_*T*(*a*(*α*),*T*(*α*))_) on the blue line to corresponding points with coordinates (*acc*_*T*(*a*(*α*),*T*(*α*))_, *trr*_*T*(*a*(*α*),*T*(*α*))_) on the orange line. Here, one should note that, in the actual validation, only the pair of thresholds (*a*(*α*), *b*(*α*)) that maximises the TRR for a target accuracy on the validation set is fitted on the independent test set *T*. However, for post-validation visualisation purposes, we showed all pairs of predictive thresholds applied to T, to demonstrate the impact of the predictive thresholds on the test performance.

### Detailed Performance Characteristics

PANProfiler Breast (ER, PR, HER2) can return a positive, negative, or indeterminate result. Positive or negative output is only given when the device is certain about its prediction. Otherwise, a prediction will not be given and the case will continue with the standard of care procedures.

To evaluate the performance of the device, confident predictions (i.e. the predictions provided by the device when it is certain) are compared to the well-characterised ground truth based on actual biomarker receptor results determined by IHC and ISH. Considering the actual and predicted results, the outcome of the device shall fall into one of the four categories of **true positive, true negative, false positive**, or **false negative**.

- **True Positive (TP)**: The device correctly predicts a positive case.
- **True Negative (TN)**: The device correctly predicts a negative case.
- **False Positive (FP)**: The device incorrectly predicts a positive case as negative.
- **False Negative (FN)**: The device incorrectly predicts a negative case as positive.

The following matrix visualises the four definitions:

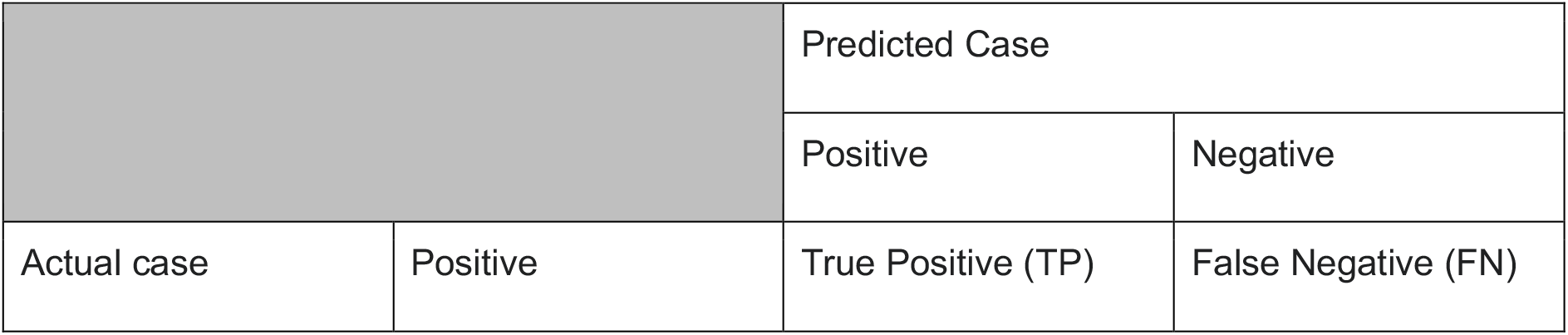
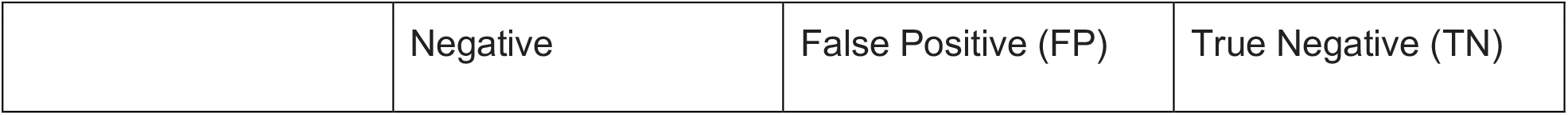

Performance characteristics used to evaluate the performance of the device are derived based on these outcomes and explained in detail in the remainder of this document.

**Accuracy**: Measures the ratio of correct predictions to all predictions done by the device.

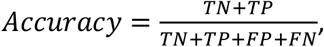

**Positive Predictive Value (PPV):** Measures the effectiveness of a device in separating true positive cases from false positive cases in the tested population. PPV is defined as follows:

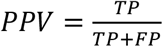

**Negative Predictive Value (NPV)**: Measures the effectiveness of a device in separating true negative cases from false negative ones in the tested population. NPV is defined as follows:

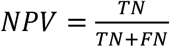

**Test Replacement Rate:** Measures the frequency of the device returning a confident prediction and is defined as the percentage of predictions done by the device over all the tested cases.

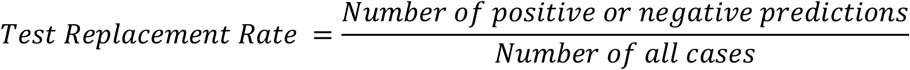

#### Area Under the Receiver Operating Characteristics Curve

A receiver operating characteristic (ROC) curve plots the relationship between TPR and FPR across different predictive thresholds (**Figure S-3**). The area under a ROC curve (AUC) describes the total area under the curve, normalised to unit square of 1. An AUC of 0.5 denotes a random model, while a perfect model that can predict all samples correctly has an AUC of 1.

**Figure S-3:**
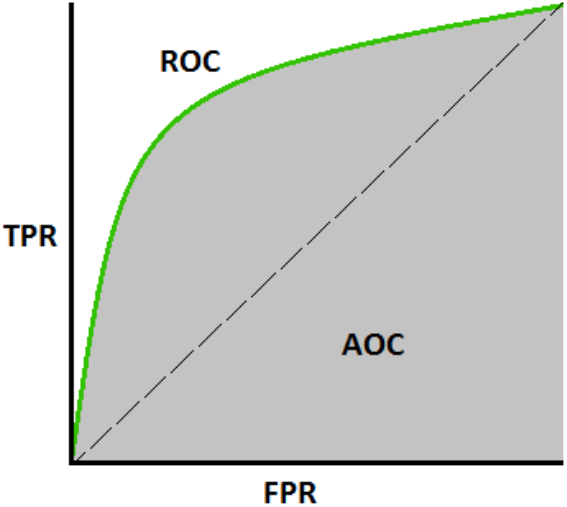
the visualisation of a ROC curve and AUC. Taken from https://bit.ly/3aFfhRz

## Supplementary Table S-1

### Laboratory Abbreviations

**Table S2:** Abbreviation of laboratories used in the confounding factor analysis figures.

| Abbr. | Laboratory Name |
| --- | --- |
| 3C | Columbia University |
| 5T | Holy Cross |
| A1 | UCSF |
| A7 | Christiana Healthcare |
| A8 | Indivumed |
| AC | International Genomics Consortium |
| AN | Cureline |
| AQ | UNC |
| B6 | Duke |
| BH | University of Pittsburgh |
| C8 | ILSBio |
| E2 | Roswell Park |
| E9 | Asterand |
| GI | ABS |
| GM | MD Anderson |
| JL | ABS |
| LD | Hartford Hospital |
| LL | Candler |
| LQ | Gundersen Lutheran Health System |
| OK | Mount Sinai School of Medicine |
| OL | University of Chicago |
| PE | Fox Chase |
| UL | Boston Medical Center |
| WT | University of Kansas |
| XX | Spectrum Health |
| Z7 | John Wayne Cancer Center |

**Table S3:**
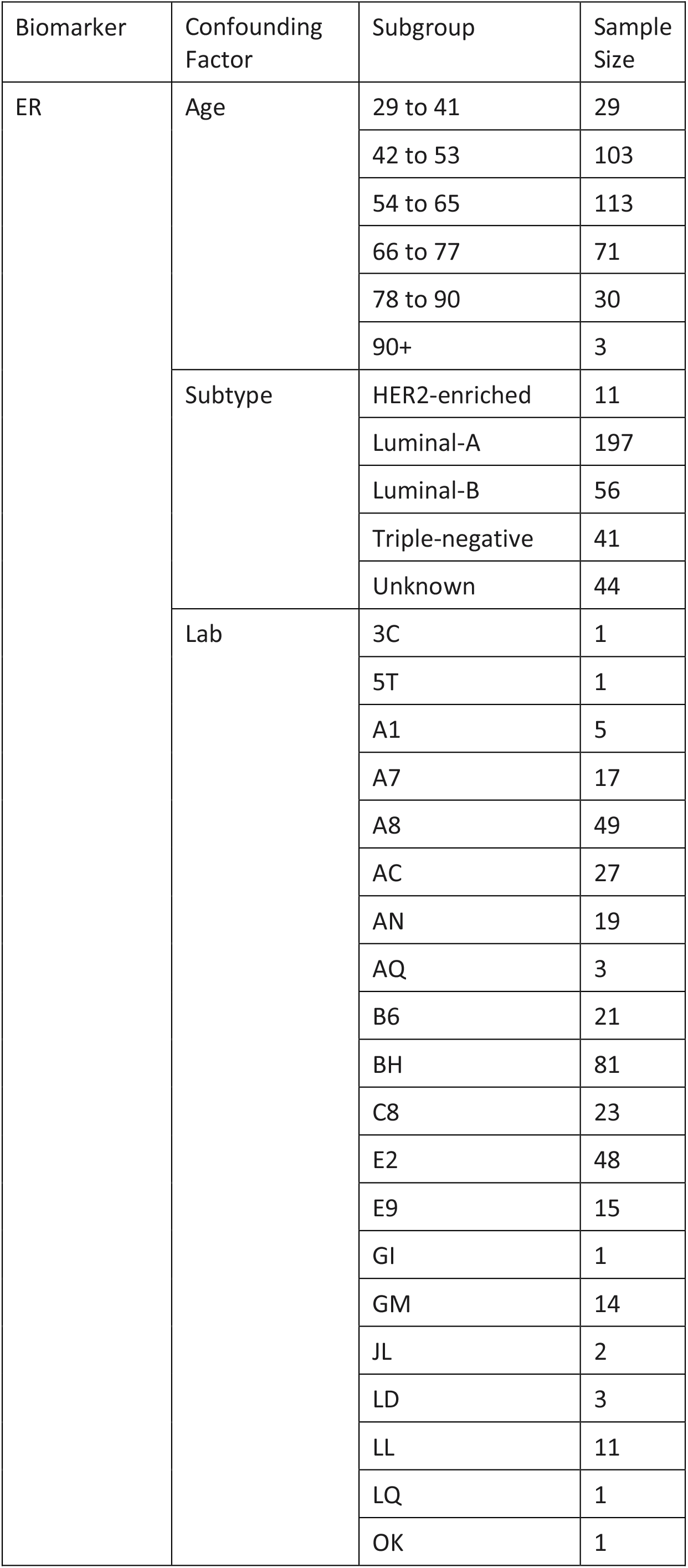
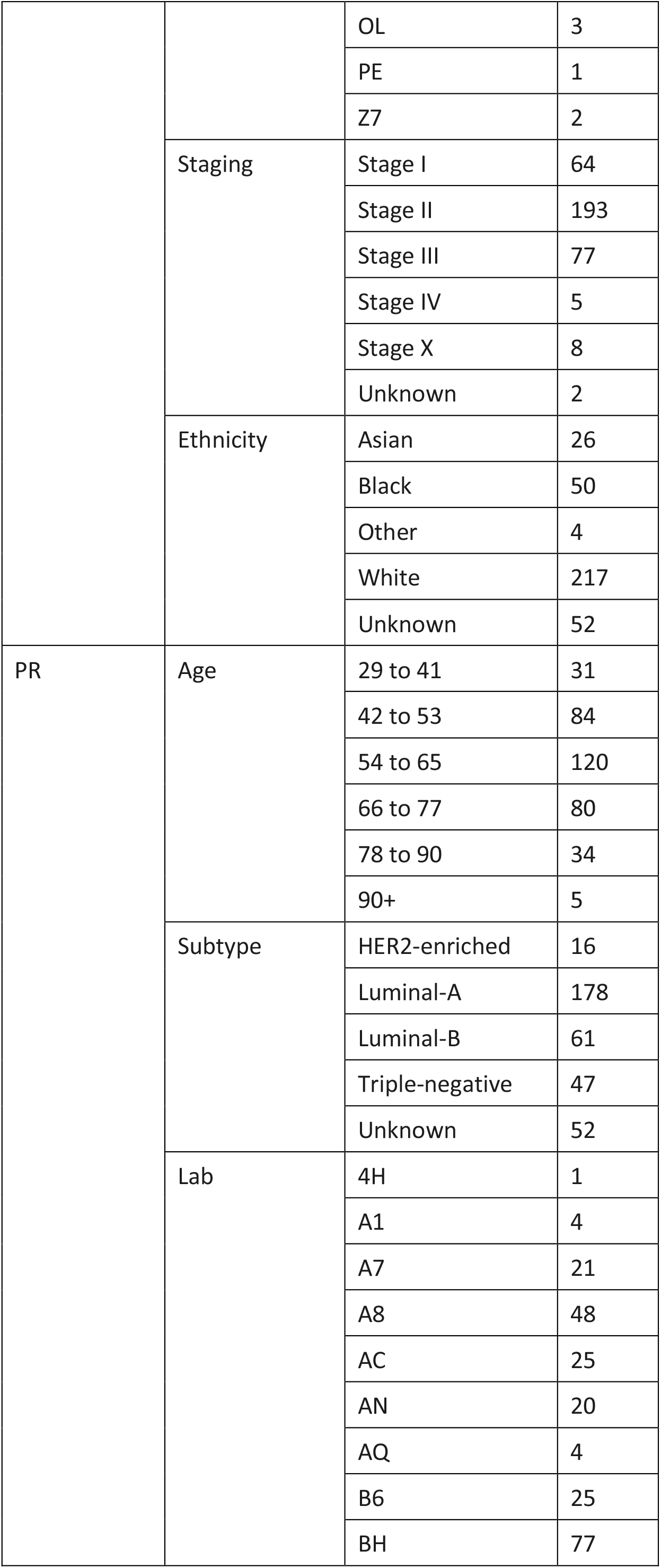
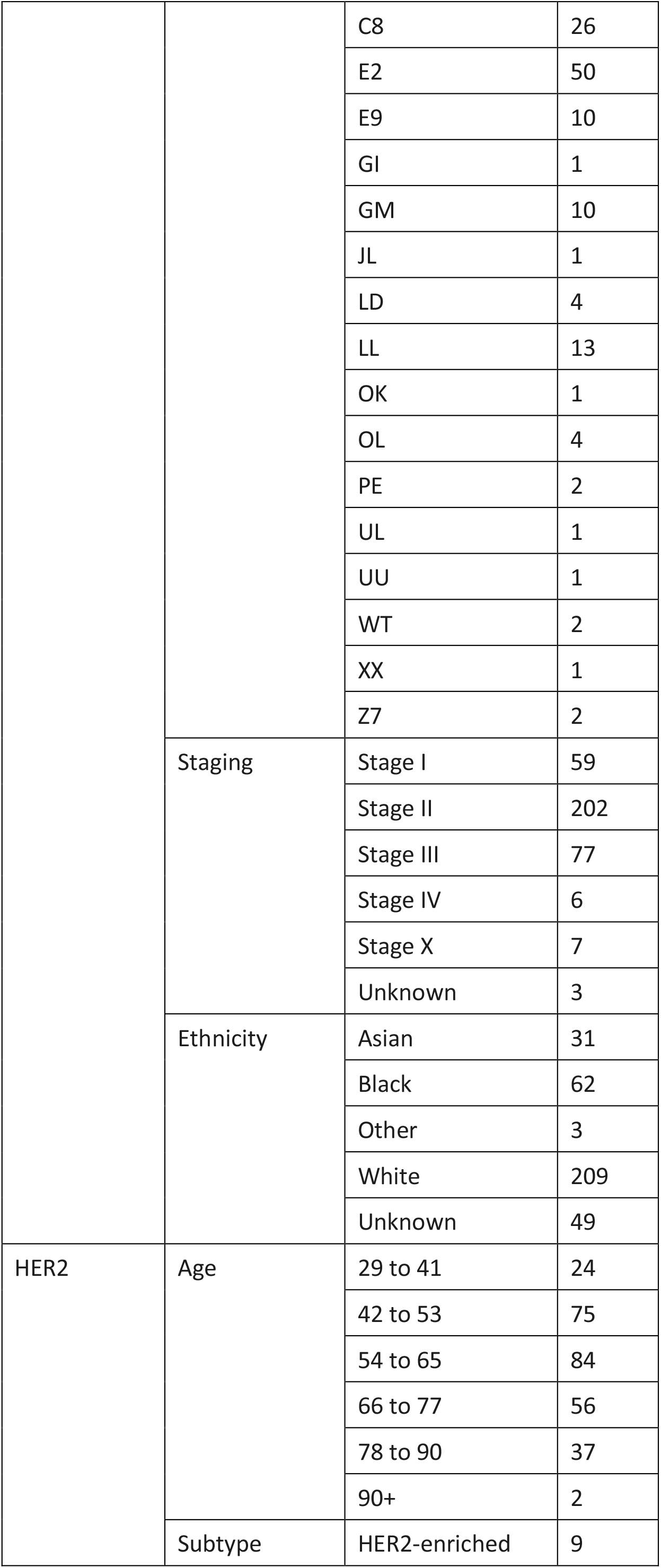
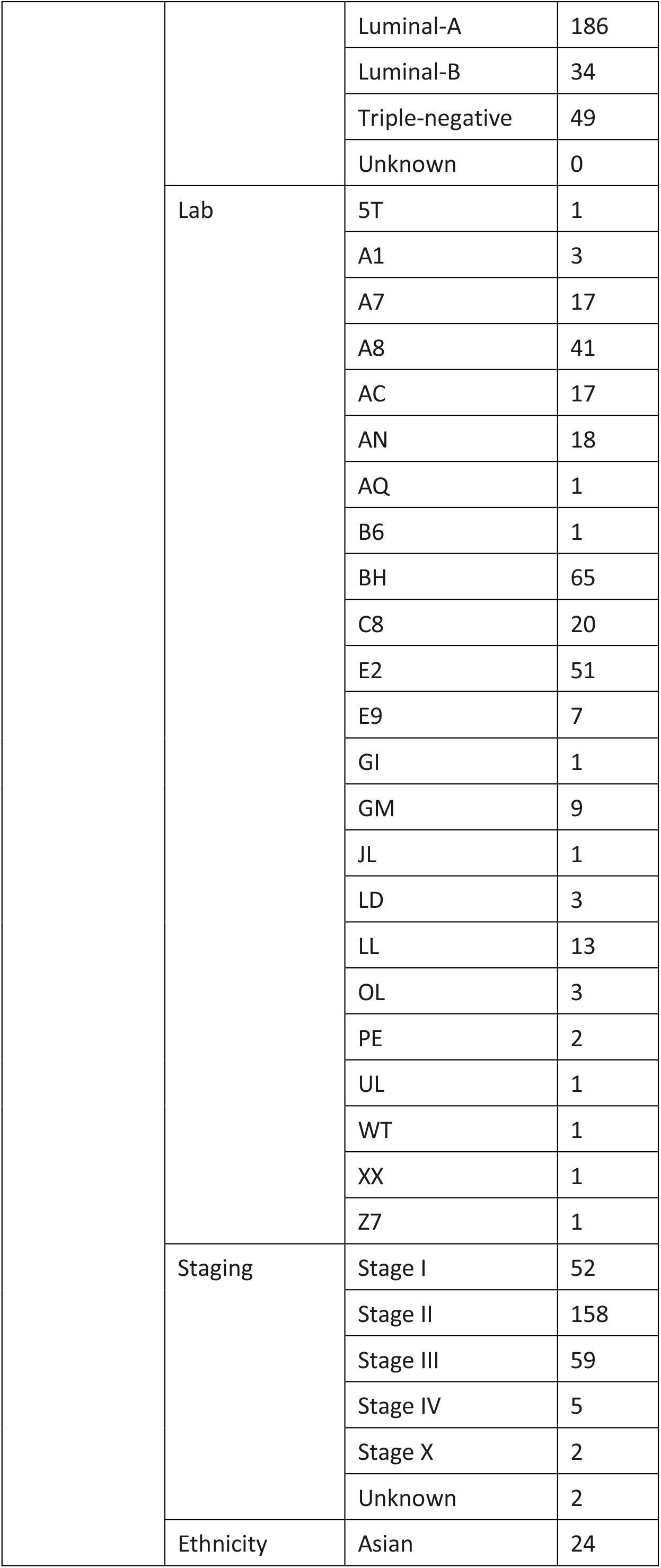
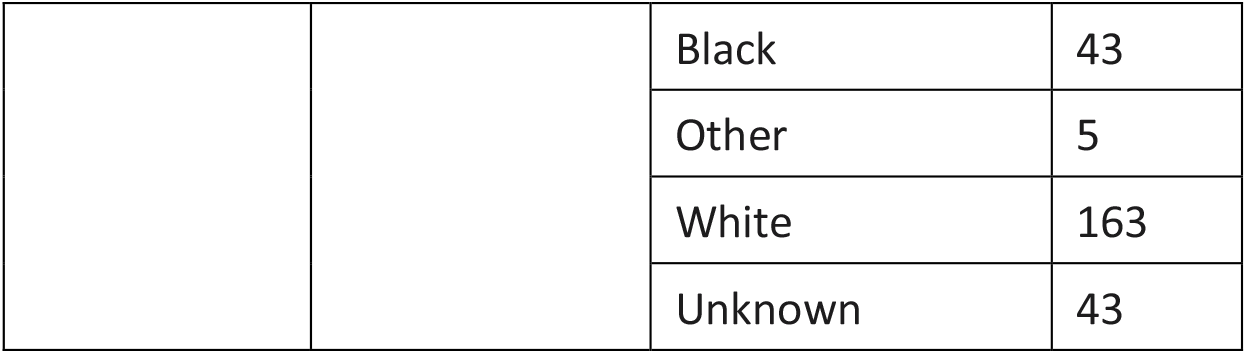
Number of cases in each subpopulation included in the confounding factor analysis.

